# PhyloNOW: Using Variable Window Sizes for Phylogenomic Analyses of Whole Genome Alignments

**DOI:** 10.64898/2026.03.04.709403

**Authors:** Jeremias Ivan, Robert Lanfear

## Abstract

Many phylogenomic studies used non-overlapping windows to address gene tree discordance across a set of aligned genomes. Recently, Ivan et al. (2025) proposed an information theoretic approach to choose an optimal window size given the alignment. However, this approach selects only a single fixed window size per chromosome, which is a useful first step but fails to account for variation in the size of non-recombining regions along each chromosome. In this study, we extend the approach of Ivan et al. (2025) and propose PhyloNOW (phylogenomic non-overlapping windows) that allows window sizes to vary across the chromosome. We show that PhyloNOW outperforms the fixed-size approach on a wide range of simulated datasets. Applying the new method on two empirical datasets from *Heliconius* butterflies and great apes, we show that window sizes vary substantially across chromosomes. Our study highlights the limitations of using a fixed window size in non-overlapping window analyses, and proposes PhyloNOW that allows for variable window sizes across whole genome alignments. PhyloNOW is available at https://github.com/jeremiasivan/PhyloNOW.

## Introduction

Many phylogenomic studies of closely-related taxa begin by aligning the genomes of species or individuals, and then use those alignments to infer evolutionary events such as speciation, introgression, and demographic changes (e.g., Pease et al. 2016; Morales-Briones et al. 2021; Lescroart et al. 2023; Herrig et al. 2024). As different sites in the alignments may reflect different evolutionary histories (Maddison 1997; Degnan and Rosenberg 2009; Mallet et al. 2016; Hibbins et al. 2023; Steenwyk et al. 2023), methods that account for this variation are important. For example, Thawornwattana et al. (2018, 2022) accounted for variation in evolutionary history across the genome by dividing individual chromosomes into blocks of 100 loci (where a locus was defined as a coding or non-coding region with a minimum length of 100bp and spaced at least 2kb apart) and inferring a species tree for each block using BPP (Flouri et al. 2018). This approach is useful for identifying introgressed regions of the genome, but excludes the majority of information contained in the genome. Additionally, assuming a fixed locus length of 100bp ignores the possibility that non-recombining loci could be substantially longer or shorter than 100bp, which may affect the accuracy of downstream inferences. To infer the complete histories of whole genome alignments – which is important for analyses such as PhyloGWAS (Pease et al. 2016) – many phylogenomic studies partition the genome alignment into non-overlapping blocks (or windows) and infer a single gene tree for each window (e.g., Fontaine et al. 2015; Pease et al. 2016; Edelman et al. 2019; Meleshko et al. 2021; Yang et al. 2021; Feng et al. 2023; Lescroart et al. 2023; Herrig et al. 2024). However, one of the main challenges for this approach is to select a window size that best reflects the recombination breakpoints of a given alignment, with a trade-off between concatenation preferring the primary tree topology (when the window size is too long, thus recombination is ignored) and gene tree estimation error (when the window size is too short, thus limiting the information available to estimate a tree for each window).

Recently, Ivan et al. (2025) proposed that the AIC (Akaike Information Criterion; Akaike 1974) – which reflects how the substitution models and the trees fit the observed alignment data – could be used to select a window size in phylogenomic non-overlapping window analyses. They showed that the AIC was a good predictor of the accuracy of different window sizes in recovering the ‘true’ histories of individual sites along simulated chromosomes. Applying this approach on empirical datasets, Ivan et al. (2025) further showed that the best window sizes for chromosome alignments of *Heliconius* butterflies ranged between ≤125-250bp, while the window sizes for great apes ranged between 500bp-1kb (and 4kb for the mitochondrial genome). These windows are significantly shorter than the ones used in many previous studies (e.g., Edelman et al. (2019) used 10kb and 50kb windows to analyse the genomes of *Heliconius* butterflies), but still longer than the estimated lengths of non-recombining blocks – sometimes called ‘coalescent genes’ or ‘*c*-genes’ – of each group. For example, Springer and Gatesy (2016) estimated the average length of *c*-genes between human, chimpanzee, and gorilla to be ∼109bp (but see also Edwards et al. (2016)). While the AIC provides a less arbitrary way to choose a window size for non-overlapping window analyses, the method proposed by Ivan et al. (2025) has two key limitations: it was evaluated using simulated chromosomes with constant recombination rates, and it selects a single fixed window size for each chromosome. A single fixed window size is problematic as some non-recombining blocks will be much longer than others even under constant recombination rate, due to the stochastic nature of recombination. And since recombination rates vary both along and between chromosomes for a variety of reasons (Jensen-Seaman et al. 2004; McVean et al. 2004; Chan et al. 2012; Edelman et al. 2019), the best window size may also vary systematically along and between chromosomes.

In this study, we extend the method proposed in Ivan et al. (2025) and propose PhyloNOW (phylogenomic non-overlapping windows) that allows the window size to vary along chromosomes using a splitting-and-merging approach (Fig. 1). We do this by iteratively splitting individual windows and/or merging neighbouring windows, and keeping such modifications when they lead to improvements in the AIC score (Fig. 1; see Methods for more details). PhyloNOW works similarly to GARD (Genetic Algorithm for Recombination Detection) (Kosakovsky Pond et al. 2006), which aims to detect recombination breakpoints from multiple sequence alignments. However, our method differs in using: (i) Maximum Likelihood (ML) instead of Neighbour-Joining (NJ) to build the gene trees; (ii) the AIC instead of the corrected AIC (*AIC*_*C*_) to score the split because both criteria yielded very similar results in prior work (Ivan et al. 2025, results not shown); and (iii) a greedy algorithm instead of a genetic algorithm to locate the putative genomic breakpoints, which allows our method to scale to whole genome alignments in a reasonable amount of time. To assess if using variable windows sizes improves the accuracy of the non-overlapping window method, we benchmark PhyloNOW on a wide range of simulated chromosomes with varying recombination rates and patterns. Finally, we apply our method to empirical datasets from *erato-sara* clade of *Heliconius* butterflies and great apes to infer the variation of window sizes along each chromosome alignment, and assess how this variation affects the distribution of gene tree topologies of each group.

**Figure 1.**
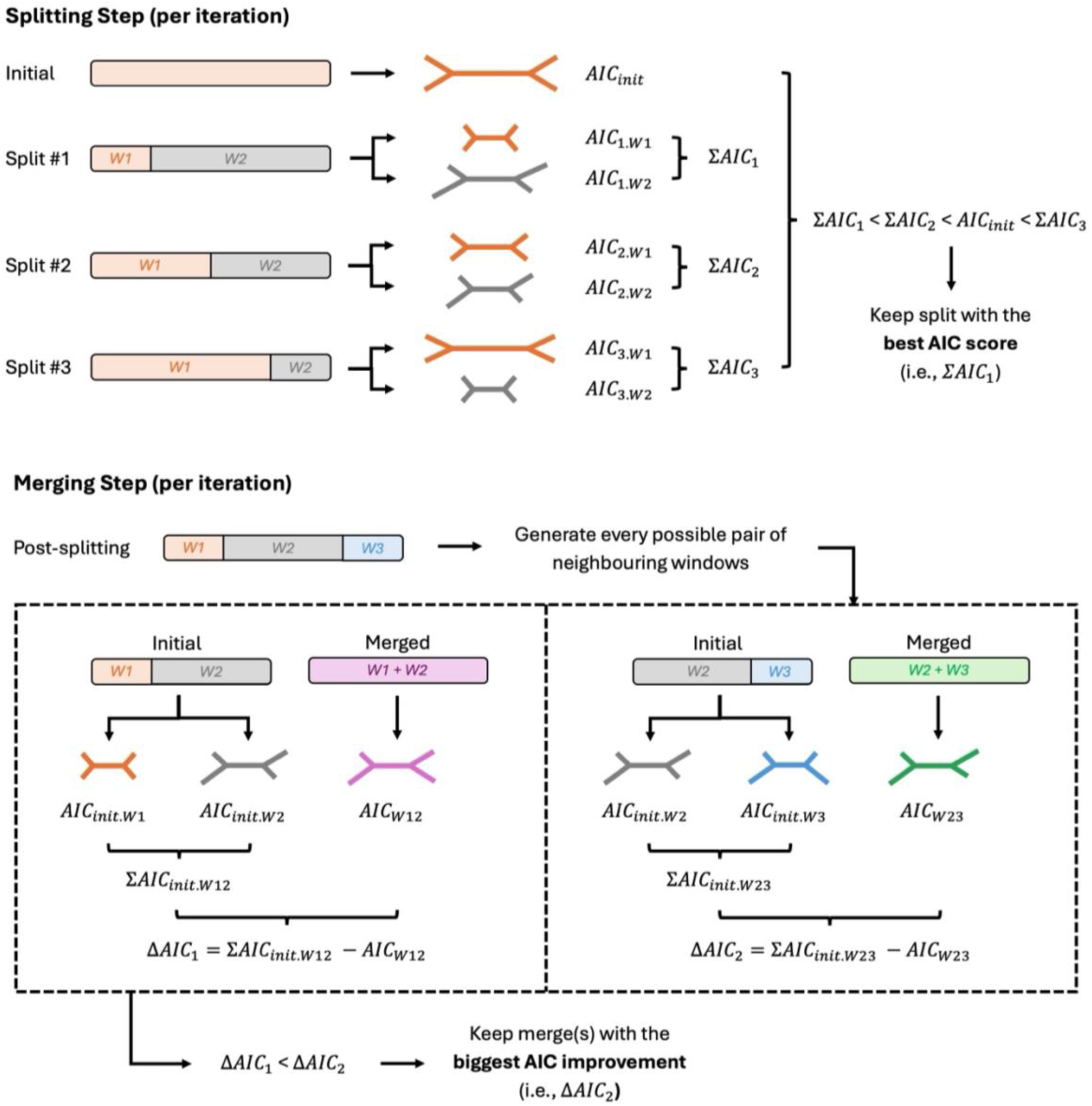
Overview of PhyloNOW approach. The splitting step is iteratively applied to every window in the alignment – selecting split configuration with the best AIC score for each window – until no further improvement can be achieved from any new split. The merging step is then iteratively applied to every possible pair of neighbouring windows in the alignment, prioritising merges with the biggest AIC improvement for each iteration until no further improvement can be achieved from any new merge.

## Materials & Methods

### Overview

PhyloNOW starts by partitioning the genome alignment into non-overlapping windows of a fixed size. Then, we proceed with the splitting step, where we attempt to split each window into two sub-windows using three size ratios (0.25:0.75, 0.50:0.50, and 0.75:0.25), building a gene tree for each window in every configuration. If the total AIC of any split configuration is lower than the AIC of the initial window, we keep the best-scoring split; otherwise, we retain the initial window. This process is repeated across all windows until no further split improves the AIC. We then enter the merging step, where for every pair of neighbouring windows, we compare the total AIC of the two initial windows and the AIC of the newly-merged window. Using a greedy algorithm, we prioritise and keep merges with the greatest AIC improvements, excluding those that do not improve the AIC and/or include a window that has already been merged in the same iteration. This is repeated until no further merge improves the AIC. The final set of windows is returned as the partitioning scheme of the genome alignment, in which the window size can be longer in some regions due to merging, and shorter in other regions due to splitting.

In order to evaluate if using variable window sizes improves the accuracy of non-overlapping window method to recover the ‘true’ histories from an alignment (compared to using only one fixed window size), we ran PhyloNOW on simulated chromosomes with varying recombination rates and patterns, and measured the accuracy of the final sets of window sizes using the two metrics used in Ivan et al. (2025): site accuracy and gene-tree root mean squared error (RMSE). Site accuracy simply records the proportion of sites in the genome that have been assigned to the correct simulated evolutionary history, while gene-tree RMSE records the RMSE between the simulated and recovered distributions of gene tree topologies. We applied our method to genome alignments from the *erato-sara* clade of *Heliconius* butterflies (Edelman et al., 2019) and great apes (Waterson et al. 2005; Locke et al. 2011; Scally et al. 2012; Schneider et al. 2017). All the code necessary to reproduce the methods in this paper is available at https://github.com/jeremiasivan/PhyloNOW.

### Simulating Chromosomes with Varying Recombination Rates

We designed seven scenarios to simulate 10Mb alignments: four with homogeneous (i.e., constant) recombination rates and three with heterogeneous recombination rates (Fig. 2). The four scenarios with homogeneous recombination rates (Fig. 2) were taken directly from Ivan et al. (2025). The three scenarios with heterogeneous recombination rates comprise: (i) *Scenario 1* (Fig. 2), a simple artificial scenario in which the first half of the 10Mb chromosome was simulated with no recombination, and the second half was simulated with an intermediate recombination rate of *ρ*=100; (ii) *Scenario 2* (Fig. 2) crudely mimics a pattern of recombination often seen in empirical datasets where the recombination rates are higher towards the ends of the chromosomes and lower towards the center, in which 2.5Mb at each end of the chromosome was simulated with *ρ*=50, and the central 5Mb region was simulated with *ρ*=0; (iii) *Scenario 3* (Fig. 2) mimics the same scenario, but with a 1Mb region at each end of the chromosome was simulated with *ρ*=200, the next 2.5Mb region at each end simulated with *ρ*=50, and the central 3Mb of the chromosome simulated with *ρ*=0.

**Figure 2.**
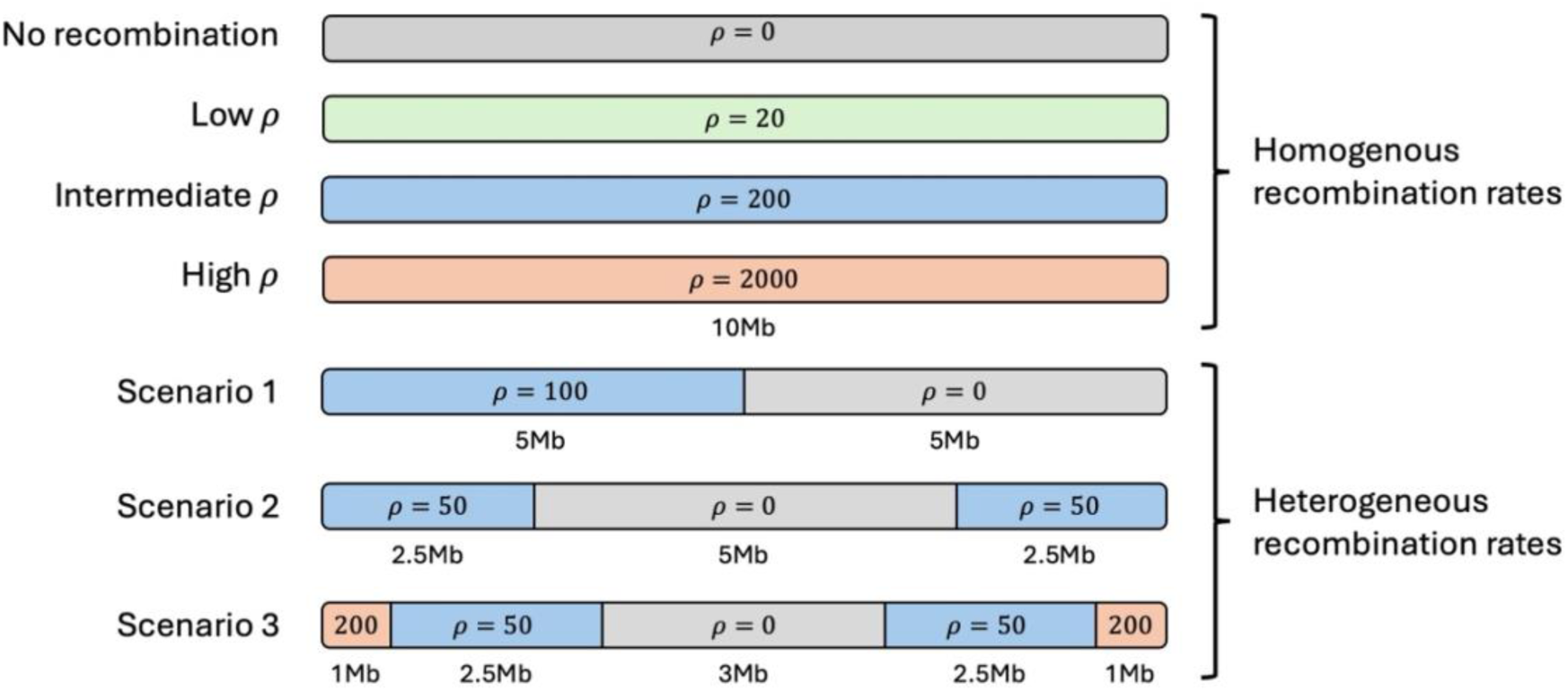
Scenarios for simulating chromosome alignments. Simulated chromosomes with homogeneous recombination rates were derived from Ivan et al. (2025), while ones with heterogeneous recombination rates were generated in this study. Gray: zero recombination rate; green: low recombination rate; blue: intermediate recombination rate; red: high recombination rate.

For each simulation, we simulated gene trees using ms (Hudson 2002) to mimic the evolutionary histories between seven *Heliconius* butterflies with a medium level of ILS, following the approach and parameters described in Ivan et al. (2025). Then, we used these sets of gene trees as input for AliSim (Ly-Trong et al. 2022) to simulate alignments under the Jukes-Cantor (JC) model (Jukes and Cantor 1969, chosen for computational efficiency). Following Ivan et al. (2025), we multiplied the coalescent branch lengths used in ms by a factor of 0.005 to produce branch lengths in substitutions per site used in AliSim. This scaling factor was shown previously to produce alignments that closely mimic empirical alignments, with roughly 5% informative sites (Ivan et al. 2025). We simulated ten independent replicates for each scenario, resulting in 70 simulated chromosomes (i.e., 10 replicates for each of the 7 simulated chromosomes depicted in Figure 2).

### Running Non-overlapping Window Analyses with Fixed Window Sizes

We ran fixed-size non-overlapping window analysis on each simulated chromosome with 16 different fixed window sizes, ranging from 100bp to 10Mb, using the same parameters used in Ivan et al. (2025). Details of the command lines used in this step are provided in the Supplementary Text. For each simulated chromosome, we then calculated the site accuracy and RMSE of each window size following Ivan et al. (2025), as well as the accuracy loss incurred when choosing a window size with the best AIC (compared to window size with the highest accuracy). To avoid unnecessary computation, we retrieved the results for the 40 simulated chromosomes with homogeneous recombination rates from Ivan et al. (2025) and did not redo these analyses.

### Running PhyloNOW on Simulated Datasets

PhyloNOW requires a starting window size to be chosen by the user. We evaluated two starting window sizes for this study: (i) the best fixed window size selected by the AIC (Ivan et al. 2025) for each simulated chromosome, and (ii) a single window comprising the entire chromosome (10Mb in this case). These relatively extreme values represent two ends of a spectrum – the former tends to be very small as it accounts for short non-recombining loci in the chromosome, while the latter is the largest possible starting value. Applying both allows us to evaluate the sensitivity of PhyloNOW to the choice of starting window size.

After a starting window size was selected, we ran PhyloNOW on each simulated chromosome. As the simulated chromosomes comprised seven taxa, we also set the minimum number of informative sites per window to be 11 – meaning that every window must contain on average one informative site for each branch on the tree. We evaluated the accuracy of the final partitioning scheme for each simulated chromosome using site accuracy and RMSE (Ivan et al. 2025), and assessed if using variable window sizes improved the two accuracy measures compared to using a single fixed window size selected by the AIC. Further details of all analyses are provided in the Supplementary Text.

### Empirical Analyses

Based on results from simulated alignments, starting PhyloNOW with full concatenation is preferable for three reasons: (i) starting with a small window size caused the algorithm to become stuck in a local optimum in one of the simulated chromosomes (Fig. 3-4); (ii) starting with full concatenation improved the recovery of the long chromosomal region with zero recombination rate as a single window (Fig. S2-S4); and (iii) it is substantially more computationally efficient, as no prior computation is required to select a starting window size. We therefore started PhyloNOW with full concatenation for analyses of empirical datasets, and suggest that application of the methods presented in this paper should proceed similarly.

**Figure 3.**
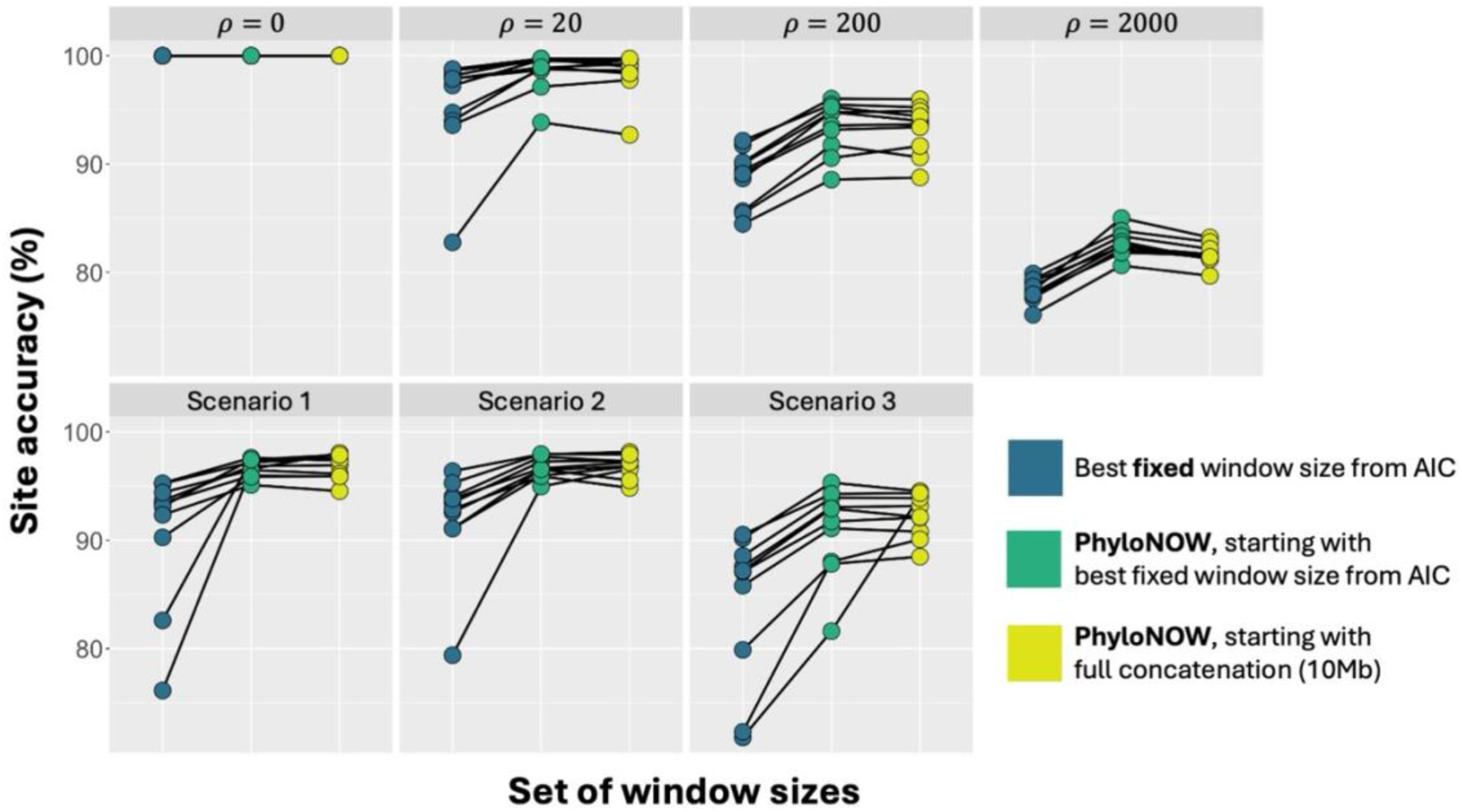
Site accuracy from running the non-overlapping window analyses with fixed and variable window sizes across 7 simulation scenarios (Fig. 2). Colour denotes the set of window sizes used when running the non-overlapping window method, either using a fixed window size (Ivan et al. 2025) or PhyloNOW with different starting window sizes. Black lines connect the same replicates.

**Figure 4.**
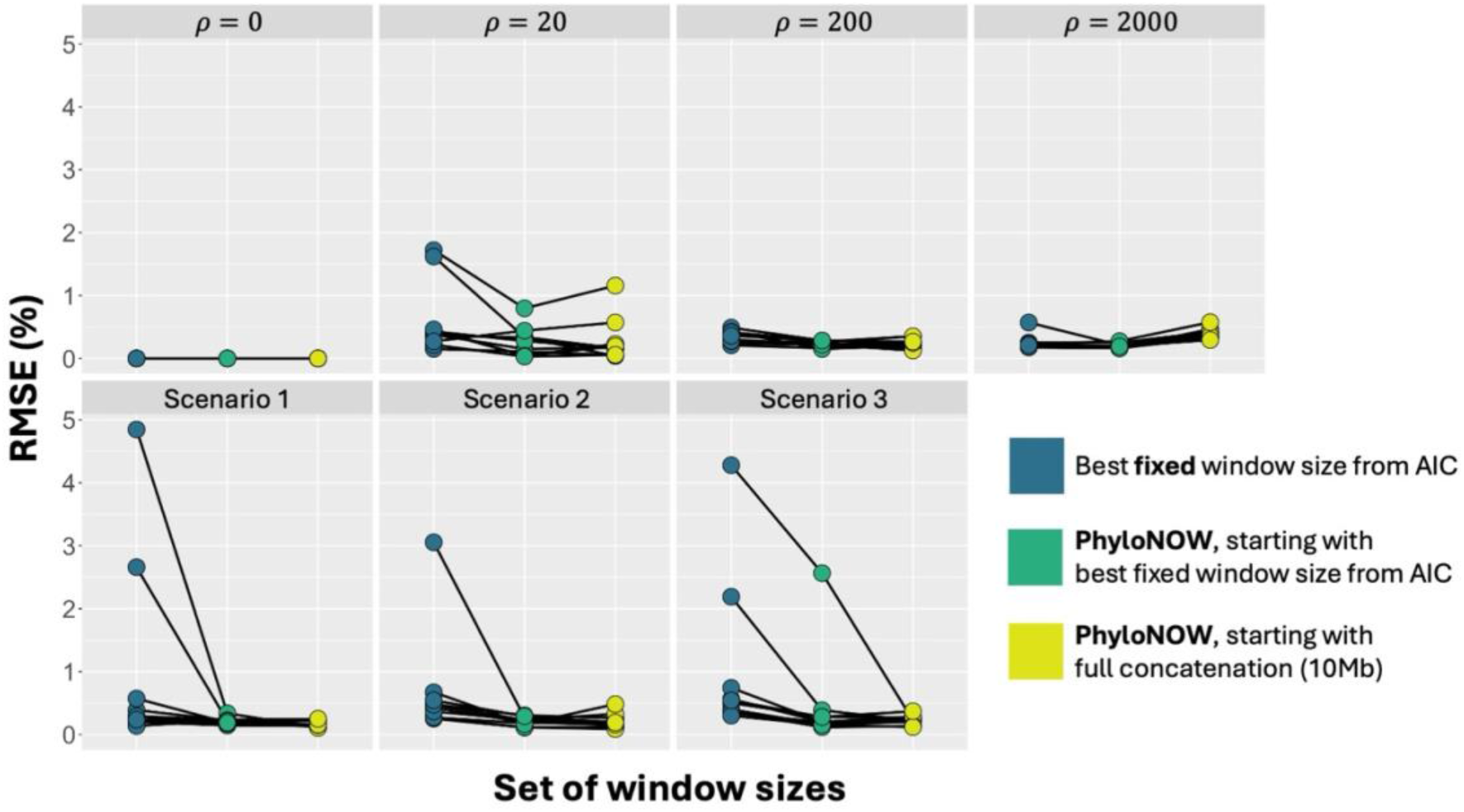
RMSE from running the non-overlapping window analyses with fixed and variable window sizes across 7 simulation scenarios (Fig. 2). Colour denotes the set of window sizes used when running the non-overlapping window method, either using a fixed window size (Ivan et al. 2025) or PhyloNOW with different starting window sizes. Black lines connect the same replicates.

#### Erato-sara clade of Heliconius butterflies

We downloaded the genome alignments from six species of *erato-sara* clade of *Heliconius* butterflies (with *H. melpomene* outgroup) from Edelman et al. (2019) following the steps described in Ivan et al. (2025). We then ran PhyloNOW on each of the 21 chromosome alignments with a minimum of 11 parsimony informative sites per window. After no further AIC improvement could be achieved from either splitting or merging, we extracted the final set of window sizes for each chromosome and rebuilt the gene tree of each window using IQ-TREE2 to estimate the bootstrap support for individual branches with the following command: `iqtree2 -s file_fasta -blmin 1/window_size -bb 1000`, where `-bb` reflects the number of UFBoot replicates (Hoang et al. 2018). We then retrieved the topology of every gene tree, and compared the proportion of each topology across the chromosomes to those from Edelman et al. (2019) and Ivan et al. (2025).

As recombination rates are generally higher on shorter chromosomes (Martin et al. 2019), we tested for a correlation between chromosome length and per-chromosome median and mean window size using R (R Core Team 2023) under three models: linear (in which window size increases proportionally with chromosome length), logarithmic (in which window size increases rapidly at first but slows down as chromosome lengthens), and asymptotic (in which window size approaches a maximum value as the chromosome length increases). We excluded chr21 (sex chromosome) from the analyses because it has distinct evolutionary history compared to autosomes (Van Belleghem et al. 2018; Martin et al. 2019), which might confound the results. All code necessary to run these analyses are available at https://github.com/jeremiasivan/PhyloNOW/blob/main/summarise_multiple_runs.R.

#### Great apes

We downloaded genome alignments of four great apes (human, chimpanzee, gorilla, and orangutan) from UCSC Genome Browser (Kent et al. 2002) following the steps described in Ivan et al. (2025). Similar to the analyses on the *Heliconius* butterflies, we ran PhyloNOW on each of the 25 chromosomes (22 autosomes, chromosome X, chromosome Y, and mitochondrial DNA) but with a minimum of 5 parsimony informative sites per window. We then built the gene trees based on the final partitioning scheme with 1,000 UFBoot replicates using IQ-TREE2 and calculated the topology distribution of the group across the chromosomes, as above. We also assessed the correlation between chromosome lengths and per-chromosome median and average window sizes under the three models mentioned above, excluding the mitochondrial DNA, chromosome X, and chromosome Y.

## Results

### Variable window sizes outperform fixed window sizes on simulated datasets

For simulated chromosomes with no recombination, non-overlapping window analyses using either fixed window sizes (from Ivan et al. 2025) or variable window sizes (from PhyloNOW) always prefer full concatenation, as expected (Fig. 3-4). Since both methods achieve 100% site accuracy in this setting, the results cannot be distinguished. However, for simulated chromosomes with non-zero recombination rates, PhyloNOW consistently outperforms the fixed window size method. When the recombination rate is homogeneous, PhyloNOW increases the site accuracy by on average 3.0-4.7% and decreases the RMSE by on average 0.05-0.35% across scenarios (except for *ρ* = 2000 with an average 0.14% increase in RMSE; Fig. 4) compared to using the best fixed window size. For simulated chromosomes with heterogeneous recombination rates, PhyloNOW increases the site accuracy by on average 4.7-8.3% and decreases the RMSE by on average 0.57-0.81% (Fig. 4) compared to using the best fixed window size. This trend is also reflected by the AIC scores, where partitioning schemes from PhyloNOW have better AIC scores than the fixed window sizes, sometimes by a substantial margin (Fig. S1).

### PhyloNOW reveals substantial length variation in window sizes

For *erato-sara* clade of *Heliconius* butterflies, PhyloNOW produces substantially different window size results compared to the fixed window size method of Ivan et al. (2025). Using PhyloNOW, the shortest window on each chromosome ranges from 18bp to 40bp, while the longest window ranges from 31kb to 106kb (Table S1; Fig. S5). Based on the distributions of window sizes across chromosomes, the best fixed window sizes selected by the stepwise non-overlapping window method (Ivan et al. 2025) are 2-5x shorter than the average window sizes from PhyloNOW, and much closer to the first quartile of the distribution than the median (Fig. S6; Table S1).

For the genomes of great apes, the shortest window ranges from 17bp to 141bp across chromosomes, while the longest window ranges from 138kb to 4.5Mb, except for chrY (chromosome Y) and mtDNA (mitochondrial DNA; Table S2; Fig. S7). For chrY, a major proportion of the alignment was not split due to missing data (Ivan et al. 2025), driving the average window size to reach almost 200kb. On the other hand, the 16kb-long mtDNA was partitioned into 8 windows, with an average window size of 2.1kb (Table S2). Compared to the best fixed window sizes from the stepwise non-overlapping window method (Ivan et al. 2025), the average window sizes from PhyloNOW are 4-11x longer (Fig. S8).

### Median window sizes are correlated to empirical chromosome lengths

The correlation between chromosome length and per-chromosome median window size of *erato-sara* clade of *Heliconius* butterflies fits a logarithmic model with *R*^2^ = 0.791 (Fig. 5 (left)). For great apes, the median window size per chromosome follows an asymptotic correlation when plotted against chromosome length with *R*^2^ = 0.531 (Fig. S5 (right)). These correlations are not significant if the median window size is replaced by the mean window size (Fig. S9), suggesting that the mean is influenced by extreme values in the window size distribution.

**Figure 5.**
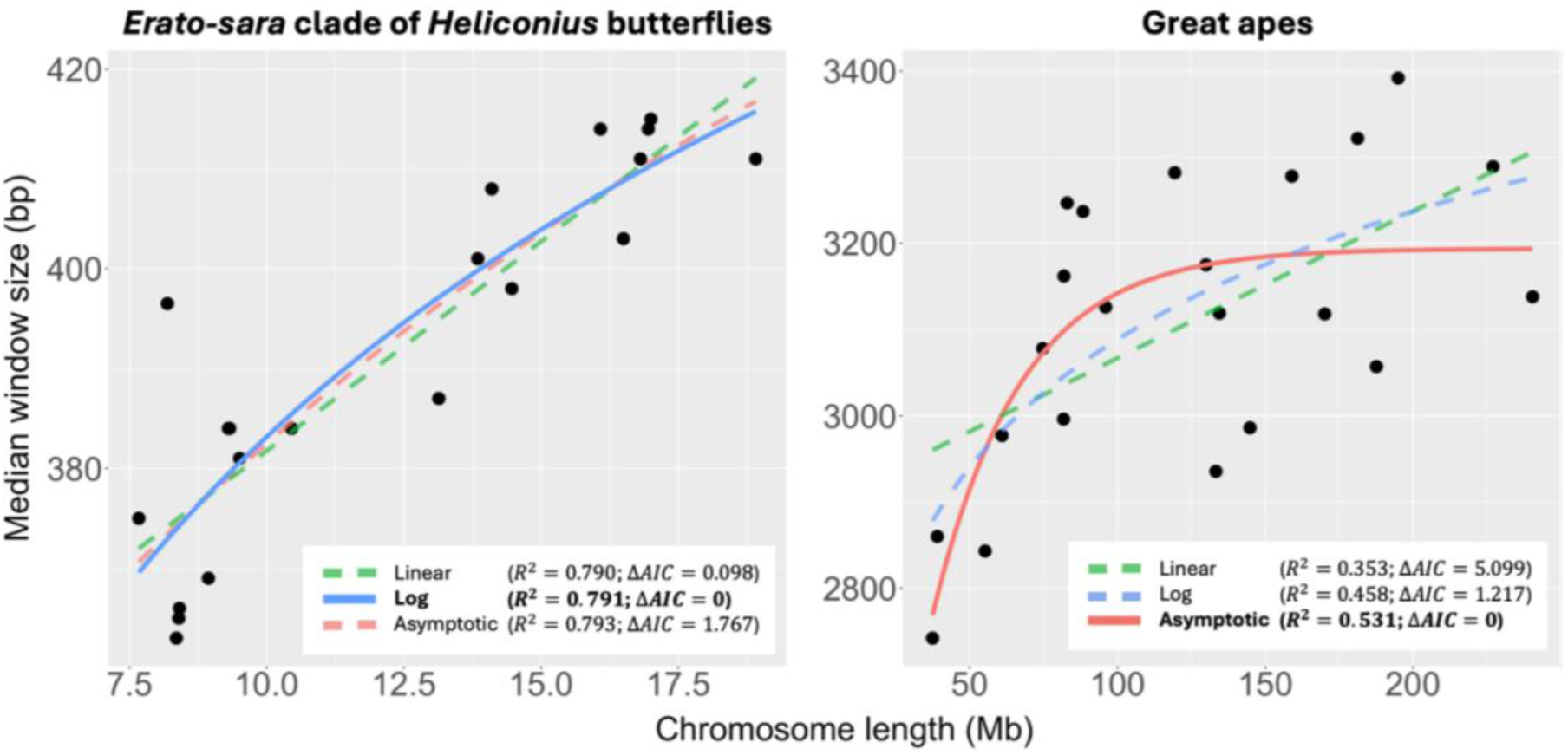
Correlation between chromosome lengths with median window sizes across autosomal chromosomes of erato-sara clade of Heliconius butterflies (left) and great apes (right). Colouring denotes different statistical models (i.e., linear, log, asymptotic). Model with the best AIC score is highlighted as solid line, while others are shown as dashed lines. Δ*AIC* reflects the AIC increase compared to the AIC of the best model.

### Window sizes can affect tree topology distribution in empirical datasets

Even though the window size varies considerably across the chromosome of *erato-sara* clade of *Heliconius* butterflies (Fig. S5-S6), the ten most common topologies recovered from the variable window size approach are consistent with those from previous studies (Edelman et al. 2019; Ivan et al. 2025), accounting for 53.0% of the sites (or 40.8% of the windows) when including all gene trees, or 97.9% of the sites (or 92.9% of the windows) from the highly-supported windows (Fig. 6, S10; Table S3-S6).

**Figure 6.**
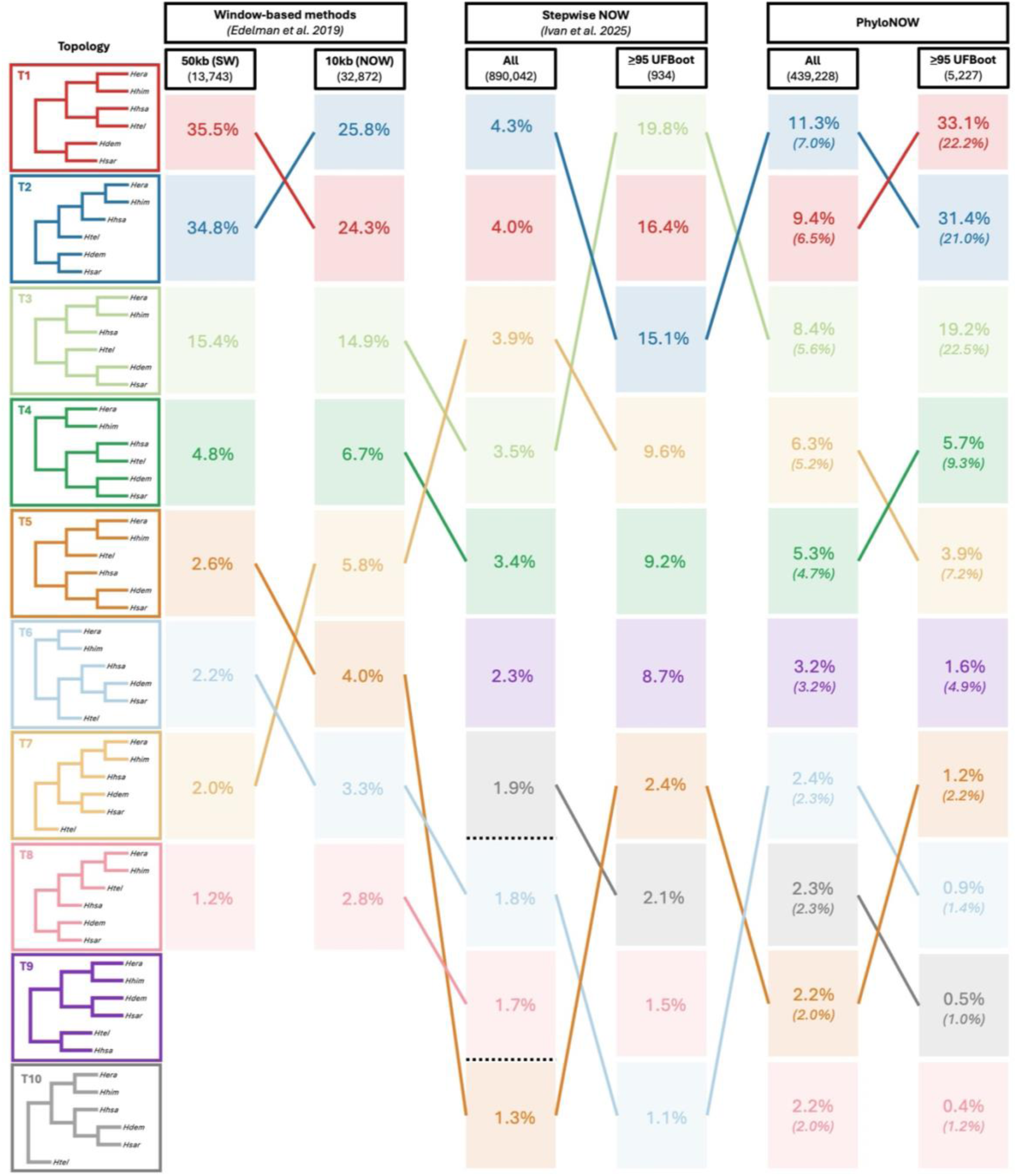
Comparison of topology frequency of the most common tree topologies from the genomes of erato-sara clade of Heliconius between 50kb sliding windows (SW) and 10kb non-overlapping windows (NOW) from Edelman et al. (2019), stepwise NOW from Ivan et al. (2025), and PhyloNOW from this study. For PhyloNOW, the percentages reflect the proportions of sites that support each topology, while the numbers in bracket reflect the proportions of windows that support each topology. Topologies are rooted with H. melpomene (not shown) as the outgroup. The number under each method reflects the total number of windows. Black dotted lines refer to other topologies not listed.

For great apes, the major topology (grouping human and chimpanzee together) is supported by 78.9% of the sites (or 75.3% of the windows), while the minor topologies account for 10.8% and 10.3% of the sites (or 12.5% and 12.2% of the windows), respectively (Fig. 7, S11; Table S7-S8). This contrasts with the topology distribution of 60.7%, 19.8%, and 19.2% when using the fixed window size approach from Ivan et al. (2025) (Fig. 7). Interestingly, two out of the eight windows from mtDNA recover each of the minor topologies (Table S7-S8; Fig. S11). This variation of gene trees in mtDNA can be either caused by systematic error (Richards et al. 2018), or associated with biological functions of the respective genomic regions (DeSalle and Tessler 2025).

**Figure 7.**
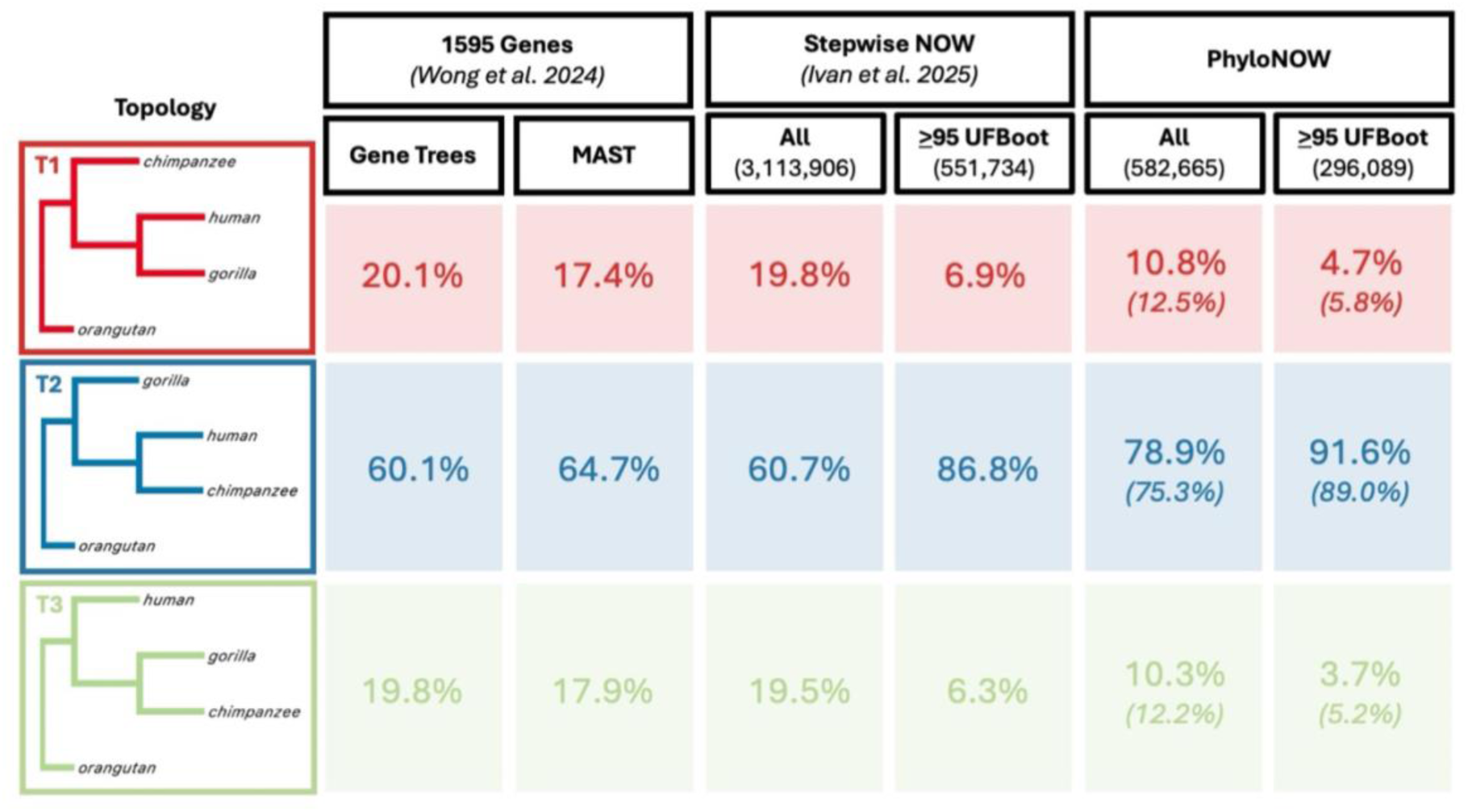
Comparison of unrooted tree topology frequency from the genomes of great apes using different approaches: single gene tree inference and MAST (Wong et al. 2024), the stepwise non-overlapping windows (NOW) that includes all gene trees and only highly-supported trees (Ivan et al. 2025), and PhyloNOW from this study. For PhyloNOW, the percentages reflect the proportions of sites that support each topology, while the numbers in bracket reflect the proportions of windows that support each topology. The number under each method reflects the total number of windows. Topologies are visualised using FigTree v1.4.4 (http://tree.bio.ed.ac.uk/software/figtree) without branch lengths.

### Variable window sizes can identify regions of biological importance

Similar to Edelman et al. (2019), we found that the proportion of T1 on the chromosomes of *Heliconius* butterflies generally goes down as the chromosome becomes longer (*R*^2^=0.743, *ρ*<0.01 for all gene trees; and *R*^2^ =0.520, *ρ*<0.01 for highly-supported trees), while T2 is positively correlated with the chromosome length (*R*^2^=0.712, *ρ*<0.01 for all gene trees; and *R*^2^=0.392, *ρ*<0.01 for highly-supported trees) (Fig. S12). When each chromosome was divided into 10 equally-sized bins along its length, we also observed that T1 has lower proportion on the ends of the chromosomes, in contrast to T2 that shows the opposite pattern (Fig. S12). Moreover, we identified two genomic regions that are dominated by the minor topologies (i.e., neither T1 nor T2; Fig. 8). The first region spans nearly 2Mb on chromosome 2 and is dominated by T3 (Fig. 8a), while the second region comprises 376 windows and is dominated by T4, covering approximately 400kb on chromosome 15 (Fig. 8b). The latter is of particular interest as it spans the loci that control the wing colour pattern in *Heliconius* butterflies (Edelman et al. 2019). To assess whether the region is best fit by a single tree topology (as PhyloNOW recovered multiple different topologies along this region despite being dominated by the T4 topology), we analysed the whole ∼400kb region as one partition, and compared the AIC to the partitioning scheme recovered from our algorithm. This analysis shows that analysing the region as a single partition has a much worse AIC score than the partitioning scheme recovered by the variable window size method (AIC score of 1,928,355 compared to 1,913,217, a difference of 15,138 units).

**Figure 8.**
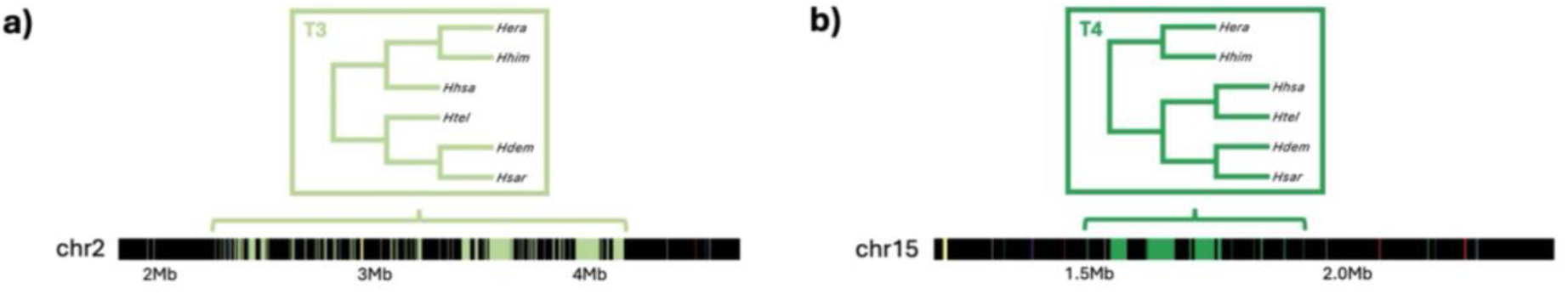
Inversion topologies on (a) chromosome 2 and (b) chromosome 15 of erato-sara clade of Heliconius butterflies (Edelman et al. 2019). Colouring is based on Fig. S6.

## Discussion

In this study, we extended the AIC-based window size selection from Ivan et al. (2025) by proposing PhyloNOW, which allows the window size to vary across the chromosome. We achieved this by iteratively splitting and merging windows until the total AIC of the whole alignment stops improving from any new split and/or merge (Fig. 1). Our results showed that PhyloNOW had better accuracy in recovering the true evolutionary history of each site in simulated datasets compared to using fixed window sizes. The fixed window sizes selected by the AIC had at least 70% site accuracy across simulated chromosomes with homogeneous and heterogeneous recombination rates (Fig. 3-4). Using variable window sizes consistently increased this accuracy further by on average 3-8% across scenarios (Fig. 3), reflecting that variable window sizes better fit the size variation of non-recombining loci across the chromosomes (Fig. S2-S4). We show that very different starting window sizes do not significantly affect the accuracy of PhyloNOW (Fig. 3-4), and therefore suggest that future phylogenomic analyses start PhyloNOW using a single window for each chromosome alignment.

When we ran PhyloNOW on the six species from *erato-sara* clade of *Heliconius* butterflies (with *H. melpomene* outgroup), we found that the window sizes varied substantially across the chromosomes (Fig. S5-S6). For instance, the window sizes on chromosome 1 ranged from 29bp to 70kb with an average of 581±11bp (95% CI), while the best fixed window size from the stepwise non-overlapping window method (Ivan et al. 2025) was 250bp (Table S1). This best fixed window size was ∼10x longer than the smallest window – which prompted the concatenation of multiple windows during gene tree inference – but 280x smaller than the biggest window – which might increase gene tree estimation error during gene tree inference. As both concatenation and gene tree estimation error might bias tree inferences (Ivan et al. 2025), this result further highlights the importance of using variable window sizes to find non-recombining loci. Unfortunately, we cannot compare the AIC between the two methods, because the stepwise non-overlapping window method (Ivan et al. 2025) excluded windows that cannot be analysed while PhyloNOW keeps them in the analysis. Interestingly, the topology distribution of the group is generally consistent across studies when considering the top ten most common topologies (Fig. 6), with either T1 or T2 as the most common topology depending on the methods being used. This might suggest that the gene tree inference for this group is quite robust to the selection of window sizes. However, we note that the *Heliconius* chromosomes from Edelman et al. (2019) were not phased. We hypothesise that phase switching in haploid representations of diploid chromosomes would lead to shorter window sizes because our approach would consider them as additional recombination breakpoints.

Similar to the results from *Heliconius* butterflies, running PhyloNOW on the genomes of great apes showed that the window sizes vary substantially across the chromosomes (Fig. S7-S8). Across the great apes chromosomes, the estimated proportion of the major topology that groups human and chimpanzee together (∼79% of the sites) was generally higher than previous estimates that ranged between 60-77% (Fig. 7; Ebersberger et al. 2007; Vanderpool et al. 2020; Wong et al. 2024; Ivan et al. 2025). There are three potential explanations for the differences in the estimates. First, both window-based and gene-based inferences are likely to be affected by trade-offs between concatenation and gene tree estimation error (Ivan et al. 2025). While our method should more accurately reflect chromosomal recombination breakpoints – and thus provides a more reliable estimate of topology distribution than gene-based or fixed-window approaches – it is not entirely free from these competing factors. For instance, we did not split a window if it cannot be analysed due to missing data or an insufficient number of parsimony informative sites (as clearly shown for chromosome Y; Table S2), which might concatenate multiple non-recombining loci together and favour the topology that is supported by the majority of sites. Second, our analyses were based on whole genome alignments that include both coding and non-coding regions. In great apes, coding regions were shown to undergo stronger selection than non-coding regions (Cagan et al. 2016). Under the Hill-Robertson effect (Hill and Robertson 1966), regions with stronger selection tend to have reduced effective population size (*N*_*e*_) (Comeron et al. 2008), which correlates with shortened coalescent time and increased gene tree discordance. As a result, a dataset that is biased towards coding regions – such as the gene-only alignments used by Wong et al. 2024 – may result in an increased proportion of discordant topologies compared to the species tree topology. However, we cannot fully evaluate this explanation because we did not explicitly differentiate the estimates between the coding and non-coding regions. Third, Reddy et al. (2017) proposed that the current substitution models might be inadequate to explain the evolution of coding regions, meaning that analyses of coding regions – on which most previous estimates were based – might underestimate the proportion of the major topology due to higher level of discordance caused by model misspecification.

Using variable window sizes, we further confirmed the findings from Edelman et al. (2019), where the proportion of T1 was higher on shorter chromosomes, while T2 was more common on longer chromosomes (Fig. S12 (left)). Martin et al. (2019) showed that chromosome length is negatively correlated with recombination rate in *Heliconius* butterflies (i.e., shorter chromosomes have higher recombination rates), which is reflected in our data by a longer median window size on longer chromosomes (Fig. 5). On the other hand, recombination rate is also correlated with the estimated admixture proportion between the three focal *Heliconius* species in that study (Martin et al. 2019). This pattern arises because genomic regions with low recombination rate tend to exhibit increased linkage among loci, which can act as a barrier to gene flow (Barton and Bengtsson 1986; Aeschbacher et al. 2017). In the *Heliconius* dataset we used here, T2 is regarded as the species tree topology while T1 reflects hybridisation between *H. hecalesia* and *H. telesiphe* (Edelman et al. 2019). As a result, T1 is enriched on shorter chromosomes with higher recombination rates and weaker barriers to hybridisation, while T2 is enriched on longer chromosomes with lower recombination rates and stronger barriers to gene flow. A similar pattern was observed along individual chromosomes (Fig. S12 (right)): T1 was more frequent towards chromosomal ends (where recombination rates are higher), while T2 had higher proportion towards the chromosomal centers (where recombination rates are lower; Martin et al. 2019). We also identified two genomic regions that are dominated by the minor topologies (Fig. 8). The first region spanned approximately 2Mb on chromosome 2 and was dominated by T3, likely caused by ILS and ancestral polymorphism (Edelman et al. 2019). The second region was 400kb-long and spanned the *cortex* locus on chromosome 15, which controls the wing colour patterns in *Heliconius* butterflies (Joron et al. 2006; Nadeau et al. 2016; Jay et al. 2018). This region reflects an inversion in *H. demeter*, *H. sara*, *H. telesiphe*, and *H. hecalesia*, but not in *H. erato*, *H. himera*, and the outgroup *H. melpomene* (Edelman et al. 2019). As a result, in this region *H. telesiphe* and *H. hecalesia* formed a sister pair, and together become the sister clade of *H. demeter* and *H. sara* (T4; Edelman et al. 2019).

Overall, PhyloNOW provides a method that uses variable window sizes to infer gene trees from whole genome alignments. The windows (i.e., loci) selected by our method should better reflect recombination breakpoints and thus serve as better approximations of *c*-genes than simply choosing a fixed window size, or selecting loci based on known genomic boundaries such as exons and introns. These loci can then be used in a number of downstream applications. For example, if the aim is to reconstruct different species trees across a genome alignment, loci from our method could be used as the starting point for coalescent-based approaches such as those used by Thawornwattana et al. (2018, 2022). If the aim is to infer the evolutionary history of every site in the genome (as may be required for applications such as PhyloGWAS (Pease et al. 2016)), then gene trees could be estimated for each locus. If stochastic error is a concern, gene trees inferred from these loci could be further refined using more complex substitution models that better-explain empirical datasets (e.g., GHOST (Crotty et al. 2020) that models heterotachy, MixtureFinder (Ren et al. 2025) that models variation of transition rates among sites), and/or more sophisticated methods that estimate the trees for individual sites (e.g., MAST (Wong et al. 2024)) or refine the gene trees in relation to each other (e.g., Espalier (Rasmussen and Guo 2023), HMMs (Siepel and Haussler 2005)). In all these cases, more accurate approximations of *c*-genes can benefit downstream analyses, and have the potential to improve our understanding of complex genomic histories.

## Supplementary Materials

Supplementary material will be available online upon the acceptance of the paper.

## Acknowledgements

We thank Minh Bui, Justin Borevitz, the members of Lanfear and Bui Lab from Australian National University, as well as Matt Hahn from Indiana University, for comments on the study design and the manuscript.

