## Supplementary Materials for "PhyloNOW: Using Variable Window Sizes for Phylogenomic Analyses of Whole Genome Alignments"

### SUPPLEMENTARY TEXT

#### Running Non-overlapping Window Analyses with Fixed Window Sizes on Simulated Chromosomes

We ran non-overlapping window analyses on each simulated chromosome with 16 different fixed window sizes, ranging from 100bp to 10Mb. Then, we built the gene tree for each window using IQ-TREE v2.4.0 (Minh et al. 2020) with the following command: `iqtree2 -S alndir -m JC -blmin 1/window_size`, where `-S alndir` refers to the directory storing all window alignments, `-m JC` sets the substitution model to be JC (Jukes and Cantor 1969), and `-blmin 1/window_size` sets the minimum branch length for all trees to represent at least one substitution per branch, penalising gene trees that were inferred from windows with limited phylogenetic signal (Ivan et al. 2025).

#### Running PhyloNOW

For each chromosome alignment, PhyloNOW starts by dividing every initial window into two sub-windows using three size ratios: 0.25:0.75, 0.50:0.50, and 0.75:0.25 (e.g., if the initial window size is 1kb and the ratio is 0.25:0.75, then the first sub-window would be 250bp while the second sub-window would be 750bp). If the initial window was not divisible by the given ratio, the length of the first sub-window was rounded up (e.g., if the initial window size is 250bp and the ratio is 0.25:0.75, then the first sub-window would be 63bp while the second sub-window would be 187bp). Next, we inferred the gene trees for the original window and each of the three split ratios using IQ-TREE2 with the following command: `iqtree2 -s file_fasta -blmin 1/window_size`, where `-s` specifies the alignment file and `-blmin` sets the minimum branch length based on the window size. We also set the minimum number of parsimony informative sites per window to be equal to the total number of branches on the (unrooted) tree, ensuring, in principle, that each branch is supported by at least one informative site. If either of the sub-windows from a split could not be analysed (e.g., due to missing data or lack of parsimony informative sites), we skipped that splitting configuration. We then compared the AIC scores of the resulting four partitioning schemes (i.e., the original unsplit window and up to three split configurations) for each window and kept the scheme with the lowest AIC for the next iteration. After applying this approach to every window in the initial alignment, we iteratively repeated the splitting procedure until no further splits led to an improvement in the AIC.

Following this, we used the AIC to guide a procedure in which neighbouring windows are iteratively merged. For each pair of neighbouring windows in the chromosome, we first calculated the change in AIC that would result from merging them into a single window. To do this, we concatenated their alignments using SeqKit v2.4.0 (Shen et al. 2016) and inferred the gene tree of the newly-merged window using IQ-TREE2 with the same parameters as above. We then extracted the AIC of the merged window and compared it with the total AIC of the two original (unmerged) windows. This procedure provided a list of neighbouring windows whose merging would improve the AIC score. However, not all these merging events were necessarily mutually compatible, because the AIC could be improved by merging the same window with the neighbouring window to the left *and* to the right of it. To resolve these conflicts, we used a greedy algorithm that prioritises merges with the biggest AIC improvements. The algorithm proceeded by merging the pair of windows with the largest AIC improvement, then removing the pair and all conflicting merges from the list of merges that could improve the AIC score. This procedure was iterated until all possible merges that improved the AIC score were exhausted. We then iteratively repeated the entire merging step until we could find no pair of neighbouring windows whose merging would improve the AIC score.

### SUPPLEMENTARY MATERIALS

**Table S1.** Summary statistics from running PhyloNOW on the genomes of *erato-sara* clade of *Heliconius* butterflies. Length: length of the chromosome alignment in base pairs (bp); #window: total number of windows from the analysis; min: minimum window size; max: maximum window size; avg: average window size; q1-q3: quartiles of window sizes; fixed: selected window size from stepwise non-overlapping window method (Ivan et al. 2025).

| chr | length (bp) | #window | window size (bp) | | | | | | |
|---|---|---|---|---|---|---|---|---|---|
| | | | min | max | avg | q1 | q2 | q3 | fixed |
| 1 | 16810322 | 28930 | 29 | 70670 | 581.069 | 273 | 411 | 649 | 250 |
| 2 | 8186463 | 13202 | 23 | 71951 | 620.093 | 237 | 396.5 | 632 | 125 |
| 3 | 9505348 | 16921 | 25 | 55696 | 561.749 | 245 | 381 | 619 | 125 |
| 4 | 8396326 | 14834 | 28 | 83020 | 566.019 | 244 | 365 | 615 | 125 |
| 5 | 8937605 | 16593 | 26 | 74782 | 538.637 | 246 | 369 | 583 | 125 |
| 6 | 13137424 | 22910 | 19 | 46598 | 573.436 | 257 | 387 | 642 | 250 |
| 7 | 13846482 | 24478 | 37 | 60848 | 565.670 | 254 | 401 | 635 | 250 |
| 8 | 8351654 | 14794 | 32 | 82579 | 564.530 | 242 | 363 | 612 | 125 |
| 9 | 7667912 | 14146 | 40 | 85296 | 542.055 | 249 | 375 | 592 | 125 |
| 10 | 18909427 | 31185 | 32 | 31162 | 606.363 | 274 | 411 | 693 | 250 |
| 11 | 10454565 | 19143 | 18 | 65415 | 546.130 | 255 | 384 | 606 | 125 |
| 12 | 16081187 | 26129 | 25 | 106003 | 615.454 | 262 | 414 | 664 | 250 |
| 13 | 16998850 | 27921 | 26 | 47271 | 608.820 | 277 | 415 | 657 | 250 |
| 14 | 8407831 | 15251 | 27 | 39456 | 551.297 | 244 | 366 | 616 | 125 |
| 15 | 9302932 | 15278 | 27 | 87314 | 608.910 | 255 | 384 | 639 | 125 |
| 16 | 9322555 | 16923 | 29 | 69134 | 550.881 | 256 | 384 | 607 | 125 |
| 17 | 14100382 | 22870 | 24 | 61965 | 616.545 | 259 | 408 | 654 | 250 |
| 18 | 16500162 | 28219 | 29 | 68827 | 584.718 | 269 | 403 | 671 | 250 |
| 19 | 16952300 | 27260 | 21 | 40364 | 621.875 | 276 | 414 | 698 | 250 |
| 20 | 14467297 | 25251 | 32 | 57284 | 572.940 | 265 | 398 | 629 | 250 |
| 21 | 13727181 | 16990 | 34 | 74227 | 807.957 | 284 | 448 | 795 | 250 |

**Table S2.** Summary statistics from running PhyloNOW on the genomes of great apes. Length: length of the chromosome alignment in base pairs (bp); #window: total number of windows from the analysis; min: minimum window size; max: maximum window size; avg: average window size; q1-q3: quartiles of window sizes; fixed: selected window size from stepwise non-overlapping window method (Ivan et al. 2025).

| chr | length (bp) | #window | window size (bp) | | | | | | |
|---|---|---|---|---|---|---|---|---|---|
| | | | min | max | avg | q1 | q2 | q3 | fixed |
| 1 | 227082584 | 45428 | 58 | 532827 | 4998.736 | 2081 | 3289 | 5550 | 1000 |
| 2 | 240480912 | 49813 | 108 | 1783351 | 4827.674 | 1959 | 3138 | 5225 | 1000 |
| 3 | 194975254 | 38612 | 41 | 203329 | 5049.603 | 2011 | 3392 | 5648 | 1000 |
| 4 | 187599630 | 40486 | 67 | 171181 | 4633.691 | 1837 | 3057 | 5159 | 1000 |
| 5 | 181263364 | 35702 | 17 | 3584554 | 5077.121 | 1969 | 3322 | 5607 | 1000 |
| 6 | 170078522 | 35396 | 87 | 1261264 | 4805.021 | 1873 | 3118 | 5261 | 1000 |
| 7 | 158966559 | 31002 | 40 | 1862890 | 5127.623 | 1942 | 3278 | 5343 | 500 |
| 8 | 144766429 | 31253 | 78 | 2544722 | 4632.081 | 1770 | 2986 | 5038 | 1000 |
| 9 | 119357962 | 22977 | 72 | 663848 | 5194.671 | 1945 | 3282 | 5511 | 1000 |
| 10 | 133258722 | 28076 | 51 | 1976432 | 4746.357 | 1832 | 2935.5 | 5146 | 500 |
| 11 | 134426162 | 26816 | 65 | 4485931 | 5012.909 | 1947 | 3119 | 5263 | 1000 |
| 12 | 129897401 | 26635 | 74 | 160548 | 4876.944 | 2009 | 3175 | 5358 | 1000 |
| 13 | 95930070 | 20694 | 97 | 145891 | 4635.647 | 1853 | 3126 | 5275 | 500 |
| 14 | 88310862 | 18179 | 63 | 327447 | 4857.850 | 1919 | 3237 | 5186.5 | 1000 |
| 15 | 81856706 | 16212 | 141 | 227636 | 5049.143 | 2001 | 3162 | 5336 | 500 |
| 16 | 81730111 | 15815 | 71 | 1818276 | 5167.886 | 1776 | 2996 | 5327 | 500 |
| 17 | 82919899 | 15507 | 72 | 2459660 | 5347.256 | 2026 | 3247 | 5406 | 500 |
| 18 | 74650691 | 16369 | 101 | 138398 | 4560.492 | 1825 | 3078 | 4866 | 500 |
| 19 | 55192149 | 13223 | 53 | 259005 | 4173.951 | 1708 | 2843 | 4797 | 500 |
| 20 | 60890884 | 12888 | 69 | 190499 | 4724.619 | 1884 | 2977 | 5291 | 500 |
| 21 | 37406299 | 8254 | 129 | 493149 | 4531.900 | 1625 | 2742 | 4628 | 500 |
| 22 | 39008215 | 8102 | 79 | 3657020 | 4814.640 | 1694 | 2860 | 4827 | 500 |
| X | 154858262 | 25087 | 75 | 4083177 | 6172.849 | 2243 | 3785 | 6728 | 1000 |
| Y | 25917497 | 131 | 10136 | 2050115 | 197843.489 | 42710.5 | 81083 | 176640 | 500 |
| M | 16568 | 8 | 55 | 10501 | 2071.000 | 417.5 | 618.5 | 1513.25 | 4000 |

**Table S3.** Distribution of the ten most common topologies across *erato-sara* clade of *Heliconius* butterflies' chromosomes based on PhyloNOW using all gene trees. Count reflects the number of windows that supports each topology. Topologies are based on Edelman et al. (2019) and Ivan et al. (2025).

| chr | topology | | | | | | | | | |
|---|---|---|---|---|---|---|---|---|---|---|
| | **T1** | **T2** | **T3** | **T4** | **T5** | **T6** | **T7** | **T8** | **T9** | **T10** |
| 1 | 1612 | 2124 | 1940 | 1306 | 734 | 802 | 1658 | 587 | 897 | 791 |
| 2 | 962 | 893 | 1103 | 552 | 157 | 162 | 584 | 269 | 403 | 204 |
| 3 | 1249 | 1083 | 699 | 839 | 351 | 398 | 792 | 424 | 539 | 395 |
| 4 | 1160 | 650 | 432 | 848 | 330 | 325 | 469 | 327 | 573 | 339 |
| 5 | 1343 | 780 | 355 | 885 | 341 | 352 | 464 | 413 | 627 | 346 |
| 6 | 1438 | 1771 | 1345 | 1022 | 394 | 510 | 1318 | 510 | 673 | 583 |
| 7 | 1699 | 1417 | 1120 | 1311 | 523 | 634 | 1190 | 500 | 934 | 626 |
| 8 | 1449 | 764 | 450 | 917 | 218 | 236 | 480 | 337 | 645 | 244 |
| 9 | 1294 | 650 | 330 | 823 | 219 | 232 | 408 | 354 | 579 | 270 |
| 10 | 1622 | 2429 | 2331 | 1166 | 638 | 781 | 1849 | 664 | 853 | 762 |
| 11 | 1454 | 953 | 655 | 1076 | 404 | 460 | 665 | 432 | 742 | 467 |
| 12 | 1591 | 1922 | 1569 | 1151 | 557 | 603 | 1532 | 577 | 797 | 620 |
| 13 | 1435 | 2518 | 2157 | 944 | 516 | 606 | 2003 | 675 | 690 | 691 |
| 14 | 1354 | 754 | 370 | 857 | 262 | 290 | 468 | 392 | 588 | 277 |
| 15 | 1176 | 708 | 506 | 1102 | 325 | 415 | 497 | 297 | 663 | 345 |
| 16 | 1337 | 845 | 497 | 942 | 355 | 377 | 507 | 396 | 639 | 362 |
| 17 | 1414 | 1636 | 1365 | 1196 | 454 | 639 | 1215 | 454 | 789 | 598 |
| 18 | 1587 | 2008 | 1615 | 1177 | 640 | 749 | 1563 | 575 | 822 | 765 |
| 19 | 1199 | 2449 | 2125 | 958 | 492 | 594 | 1821 | 616 | 674 | 655 |
| 20 | 1686 | 1670 | 1355 | 1240 | 557 | 622 | 1268 | 591 | 874 | 611 |
| 21 | 493 | 2611 | 2154 | 272 | 135 | 178 | 2071 | 287 | 188 | 254 |

**Table S4.** Weighted distribution of the ten most common topologies across *erato-sara* clade of *Heliconius* butterflies' chromosomes based on PhyloNOW using all gene trees. Count reflects the number of sites that supports each topology. Topologies are based on Edelman et al. (2019) and Ivan et al. (2025).

| chr | topology | | | | | | | | | |
|---|---|---|---|---|---|---|---|---|---|---|
| | **T1** | **T2** | **T3** | **T4** | **T5** | **T6** | **T7** | **T8** | **T9** | **T10** |
| 1 | 1269986 | 2014516 | 1732242 | 813608 | 478139 | 514005 | 1074251 | 337534 | 522263 | 481314 |

| | | | | | | | | | | |
|---|---|---|---|---|---|---|---|---|---|---|
| 2 | 831510 | 654115 | 1280688 | 318905 | 107594 | 75633 | 356569 | 190576 | 251045 | 121489 |
| 3 | 1067081 | 995261 | 499655 | 544541 | 230306 | 228276 | 545320 | 227842 | 323564 | 249804 |
| 4 | 1095165 | 635042 | 319506 | 570728 | 199896 | 196884 | 311336 | 176381 | 343496 | 212614 |
| 5 | 1288729 | 625731 | 232333 | 589614 | 224326 | 203698 | 263871 | 238518 | 341657 | 197984 |
| 6 | 1125162 | 1613719 | 1084463 | 626441 | 263337 | 305943 | 884353 | 320518 | 370946 | 337237 |
| 7 | 1474912 | 1224607 | 870696 | 858397 | 340106 | 468256 | 795383 | 261733 | 544375 | 364046 |
| 8 | 1501586 | 585517 | 312346 | 597132 | 120454 | 138291 | 278976 | 182940 | 383435 | 128126 |
| 9 | 1338822 | 431526 | 219192 | 525900 | 143533 | 136432 | 225129 | 194572 | 353646 | 143570 |
| 10 | 1192721 | 2173366 | 2120462 | 792092 | 448734 | 550489 | 1425273 | 391114 | 507366 | 473556 |
| 11 | 1222016 | 688137 | 490217 | 715994 | 262035 | 274593 | 440024 | 256361 | 422847 | 290477 |
| 12 | 1299748 | 2122151 | 1372960 | 744057 | 393211 | 390361 | 1170995 | 345282 | 484889 | 386170 |
| 13 | 1028239 | 2679495 | 1893141 | 607044 | 338610 | 359507 | 1432575 | 394026 | 368554 | 393655 |
| 14 | 1297969 | 522704 | 262522 | 535956 | 172472 | 167507 | 280516 | 236947 | 343587 | 154847 |
| 15 | 1005681 | 654100 | 420805 | 977903 | 220598 | 244868 | 468982 | 189973 | 402662 | 192333 |
| 16 | 1237966 | 645409 | 368270 | 583670 | 216972 | 217518 | 328706 | 240748 | 408167 | 207366 |
| 17 | 1348002 | 1540480 | 1256290 | 863611 | 351458 | 427379 | 865562 | 277618 | 442995 | 368809 |
| 18 | 1225913 | 1831940 | 1440073 | 798906 | 408561 | 472622 | 1161505 | 326506 | 495211 | 456801 |
| 19 | 837698 | 2548045 | 1983014 | 624992 | 322881 | 368221 | 1374324 | 383136 | 382861 | 386872 |
| 20 | 1322830 | 1477952 | 1077207 | 839020 | 353679 | 447316 | 864536 | 330708 | 536084 | 375577 |
| 21 | 292810 | 3685505 | 2531635 | 145551 | 80175 | 108678 | 1806098 | 164268 | 110867 | 129400 |

**Table S5.** Distribution of the ten most common topologies across *erato-sara* clade of *Heliconius* butterflies' chromosomes based on PhyloNOW using gene trees with ≥95 average UFBoot support. Count reflects the number of windows that supports each topology. Topologies are based on Edelman et al. (2019) and Ivan et al. (2025).

| **chr** | **topology** | | | | | | | | | |
|---|---|---|---|---|---|---|---|---|---|---|
| | **T1** | **T2** | **T3** | **T4** | **T5** | **T6** | **T7** | **T8** | **T9** | **T10** |
| 1 | 60 | 83 | 109 | 41 | 12 | 6 | 26 | 2 | 15 | 2 |
| 2 | 33 | 21 | 133 | 8 | 2 | 2 | 7 | 3 | 5 | 0 |
| 3 | 45 | 28 | 15 | 17 | 0 | 0 | 10 | 5 | 10 | 1 |
| 4 | 42 | 13 | 12 | 14 | 2 | 1 | 2 | 1 | 10 | 1 |
| 5 | 53 | 15 | 2 | 21 | 5 | 1 | 6 | 0 | 6 | 1 |
| 6 | 63 | 63 | 51 | 29 | 7 | 5 | 18 | 4 | 12 | 2 |
| 7 | 90 | 37 | 53 | 35 | 10 | 7 | 14 | 1 | 17 | 5 |
| 8 | 71 | 17 | 8 | 25 | 7 | 1 | 4 | 1 | 8 | 1 |
| 9 | 62 | 6 | 5 | 11 | 1 | 0 | 4 | 1 | 10 | 1 |
| 10 | 66 | 85 | 137 | 28 | 7 | 2 | 36 | 5 | 18 | 4 |

| | | | | | | | | | | |
|---|---|---|---|---|---|---|---|---|---|---|
| 11 | 75 | 23 | 12 | 23 | 7 | 2 | 6 | 1 | 14 | 4 |
| 12 | 62 | 84 | 75 | 22 | 4 | 8 | 28 | 4 | 16 | 5 |
| 13 | 58 | 126 | 131 | 16 | 6 | 4 | 39 | 6 | 7 | 4 |
| 14 | 64 | 7 | 10 | 22 | 3 | 0 | 3 | 1 | 8 | 1 |
| 15 | 44 | 12 | 12 | 37 | 3 | 4 | 5 | 1 | 15 | 1 |
| 16 | 56 | 18 | 10 | 21 | 5 | 3 | 10 | 1 | 13 | 2 |
| 17 | 55 | 71 | 75 | 35 | 4 | 7 | 25 | 2 | 19 | 3 |
| 18 | 53 | 61 | 67 | 25 | 13 | 5 | 31 | 4 | 12 | 8 |
| 19 | 40 | 114 | 121 | 28 | 6 | 7 | 31 | 8 | 17 | 3 |
| 20 | 59 | 57 | 46 | 30 | 9 | 8 | 18 | 6 | 22 | 3 |
| 21 | 7 | 158 | 90 | 0 | 1 | 1 | 53 | 3 | 0 | 2 |

**Table S6.** Weighted distribution of the ten most common topologies across *erato-sara* clade of *Heliconius* butterflies' chromosomes based on PhyloNOW using gene trees with ≥95 average UFBoot support. Count reflects the number of sites that supports each topology. Topologies are based on Edelman et al. (2019) and Ivan et al. (2025).

| **chr** | **topology** | | | | | | | | | |
|---|---|---|---|---|---|---|---|---|---|---|
| | **T1** | **T2** | **T3** | **T4** | **T5** | **T6** | **T7** | **T8** | **T9** | **T10** |
| 1 | 295035 | 366651 | 317684 | 53292 | 11717 | 4915 | 46848 | 7082 | 14870 | 2273 |
| 2 | 161386 | 54206 | 627158 | 7486 | 1142 | 867 | 7246 | 4721 | 3808 | 0 |
| 3 | 175349 | 161105 | 27856 | 50723 | 0 | 0 | 44122 | 6171 | 5192 | 4182 |
| 4 | 356151 | 168642 | 40309 | 19125 | 2311 | 648 | 7662 | 585 | 10490 | 973 |
| 5 | 411676 | 32889 | 2627 | 49167 | 14503 | 1966 | 7318 | 0 | 5481 | 738 |
| 6 | 160148 | 379730 | 124642 | 42842 | 16045 | 7354 | 17846 | 4379 | 13070 | 7052 |
| 7 | 418821 | 118168 | 113742 | 60768 | 14425 | 55425 | 18228 | 1996 | 19944 | 9738 |
| 8 | 536358 | 97109 | 11482 | 41287 | 5804 | 646 | 5293 | 242 | 8719 | 612 |
| 9 | 461791 | 11023 | 7404 | 13436 | 14808 | 0 | 5386 | 250 | 14229 | 395 |
| 10 | 197318 | 291078 | 338926 | 47723 | 11916 | 5767 | 81272 | 2571 | 15579 | 9166 |
| 11 | 237308 | 70949 | 16781 | 24637 | 11875 | 2251 | 7183 | 728 | 12022 | 7978 |
| 12 | 252254 | 585087 | 154045 | 40735 | 25179 | 13656 | 56955 | 10971 | 23085 | 5917 |
| 13 | 163684 | 704468 | 280802 | 26278 | 19094 | 6925 | 65595 | 6008 | 4556 | 5403 |
| 14 | 338742 | 16240 | 14361 | 38539 | 4418 | 0 | 2685 | 1516 | 19617 | 548 |
| 15 | 200224 | 136966 | 18314 | 206435 | 5244 | 8326 | 7087 | 682 | 17652 | 455 |
| 16 | 351201 | 83360 | 22128 | 32395 | 6866 | 5401 | 11530 | 385 | 26087 | 2888 |

| | | | | | | | | | | |
|---|---|---|---|---|---|---|---|---|---|---|
| 17 | 439105 | 347007 | 244596 | 104213 | 10773 | 10432 | 33413 | 1104 | 16768 | 3769 |
| 18 | 246811 | 167826 | 226745 | 41142 | 14545 | 7929 | 60424 | 5437 | 12669 | 8814 |
| 19 | 137765 | 564297 | 316432 | 38256 | 9374 | 6037 | 75098 | 8067 | 19364 | 5439 |
| 20 | 271055 | 241002 | 84782 | 59556 | 16292 | 12394 | 28058 | 6033 | 18628 | 2625 |
| 21 | 11008 | 934386 | 381952 | 0 | 1061 | 1193 | 93257 | 4141 | 0 | 2030 |

**Table S7.** Distribution of the three unrooted topologies across great apes' chromosome based on PhyloNOW using all gene trees (left) and gene trees with ≥95 average UFBoot support (right). Count reflects the number of windows that supports each topology. H: human; O: orangutan; G: gorilla; C: chimpanzee.

| **chr** | **all gene trees** | | | **gene trees ≥95 UFBoot support** | | |
|---|---|---|---|---|---|---|
| | **[H,C\|O,G]** | **[H,G\|O,C]** | **[H,O\|C,G]** | **[H,C\|O,G]** | **[H,G\|O,C]** | **[H,O\|C,G]** |
| 1 | 35109 | 5298 | 5021 | 21423 | 1235 | 1093 |
| 2 | 37782 | 6068 | 5963 | 22680 | 1325 | 1221 |
| 3 | 28983 | 4769 | 4860 | 17628 | 1141 | 1113 |
| 4 | 28862 | 5776 | 5848 | 17367 | 1540 | 1539 |
| 5 | 26421 | 4523 | 4758 | 16070 | 1037 | 1057 |
| 6 | 26129 | 4560 | 4707 | 15656 | 1071 | 1123 |
| 7 | 23232 | 3977 | 3793 | 13772 | 968 | 748 |
| 8 | 23040 | 4151 | 4062 | 13845 | 1030 | 914 |
| 9 | 17745 | 2595 | 2637 | 10922 | 529 | 575 |
| 10 | 21011 | 3571 | 3494 | 12437 | 821 | 697 |
| 11 | 20451 | 3207 | 3158 | 12513 | 796 | 709 |
| 12 | 20205 | 3220 | 3210 | 12167 | 806 | 706 |
| 13 | 15035 | 2887 | 2772 | 8796 | 729 | 659 |
| 14 | 13666 | 2336 | 2177 | 8290 | 573 | 453 |
| 15 | 12108 | 2110 | 1994 | 6944 | 484 | 365 |
| 16 | 12173 | 1843 | 1799 | 7169 | 406 | 315 |
| 17 | 11883 | 1769 | 1855 | 6279 | 254 | 288 |
| 18 | 11757 | 2351 | 2261 | 6720 | 608 | 485 |
| 19 | 9333 | 2011 | 1879 | 4592 | 480 | 334 |
| 20 | 9709 | 1651 | 1528 | 5628 | 338 | 274 |
| 21 | 5766 | 1282 | 1206 | 3151 | 392 | 246 |

| | | | | | | |
|---|---|---|---|---|---|---|
| 22 | 5990 | 1136 | 976 | 3199 | 295 | 159 |
| X | 22379 | 1477 | 1231 | 16184 | 404 | 219 |
| Y | 120 | 4 | 7 | 99 | 2 | 1 |
| M | 6 | 1 | 1 | 1 | 0 | 0 |

**Table S8.** Weighted distribution of the three unrooted topologies across great apes' chromosome based on PhyloNOW using all gene trees (left) and gene trees with ≥95 average UFBoot support (right). Count reflects the number of sites that supports each topology. H: human; O: orangutan; G: gorilla; C: chimpanzee.

| **chr** | **all gene trees** | | | **gene trees ≥95 UFBoot support** | | |
|---|---|---|---|---|---|---|
| | **[H,C\|O,G]** | **[H,G\|O,C]** | **[H,O\|C,G]** | **[H,C\|O,G]** | **[H,G\|O,C]** | **[H,O\|C,G]** |
| 1 | 181523480 | 23530157 | 22028947 | 124074989 | 5688122 | 4578964 |
| 2 | 190247126 | 25775234 | 24458552 | 128689232 | 5793855 | 4784447 |
| 3 | 153079142 | 20645838 | 21250274 | 103996451 | 4800633 | 4723697 |
| 4 | 138758957 | 24214469 | 24626204 | 91681494 | 6028618 | 6226692 |
| 5 | 140398544 | 20581649 | 20283171 | 96344910 | 5147738 | 4125235 |
| 6 | 131956677 | 18875760 | 19246085 | 88690616 | 4448747 | 4388311 |
| 7 | 124181328 | 18582251 | 16202980 | 84295811 | 4031081 | 3461987 |
| 8 | 108922327 | 19126308 | 16717794 | 72897967 | 6619060 | 3738640 |
| 9 | 95123220 | 12295898 | 11938844 | 65970920 | 3109566 | 3161303 |
| 10 | 106087132 | 13621806 | 13549784 | 71872764 | 3038935 | 2741690 |
| 11 | 107950538 | 13532190 | 12943434 | 75271874 | 3561417 | 2850951 |
| 12 | 103454880 | 13350666 | 13091855 | 69444685 | 3071714 | 2875661 |
| 13 | 72930629 | 11710373 | 11289068 | 48185921 | 2793627 | 2535406 |
| 14 | 69615599 | 9792560 | 8902703 | 47070655 | 2123803 | 1672789 |
| 15 | 63823281 | 9544023 | 8489402 | 41709774 | 2652122 | 1526717 |
| 16 | 65753011 | 8528181 | 7448919 | 44548181 | 2017465 | 1237389 |
| 17 | 65255779 | 9583521 | 8080599 | 40433880 | 3408553 | 1368742 |
| 18 | 56889428 | 9168285 | 8592978 | 37574844 | 2336004 | 1651127 |
| 19 | 40605934 | 7472182 | 7114033 | 21748821 | 1751406 | 1191436 |
| 20 | 48122798 | 6737025 | 6031061 | 31822646 | 1742815 | 1058032 |
| 21 | 27063959 | 5933629 | 4408711 | 17058166 | 2618883 | 841826 |

| | | | | | | |
|---|---|---|---|---|---|---|
| 22 | 30656967 | 4695464 | 3655784 | 19935092 | 1525630 | 774374 |
| X | 142009632 | 6679456 | 6169174 | 112894059 | 1413193 | 1028720 |
| Y | 23228247 | 188945 | 2500305 | 19214466 | 80082 | 384395 |
| M | 13267 | 194 | 3107 | 10501 | 0 | 0 |

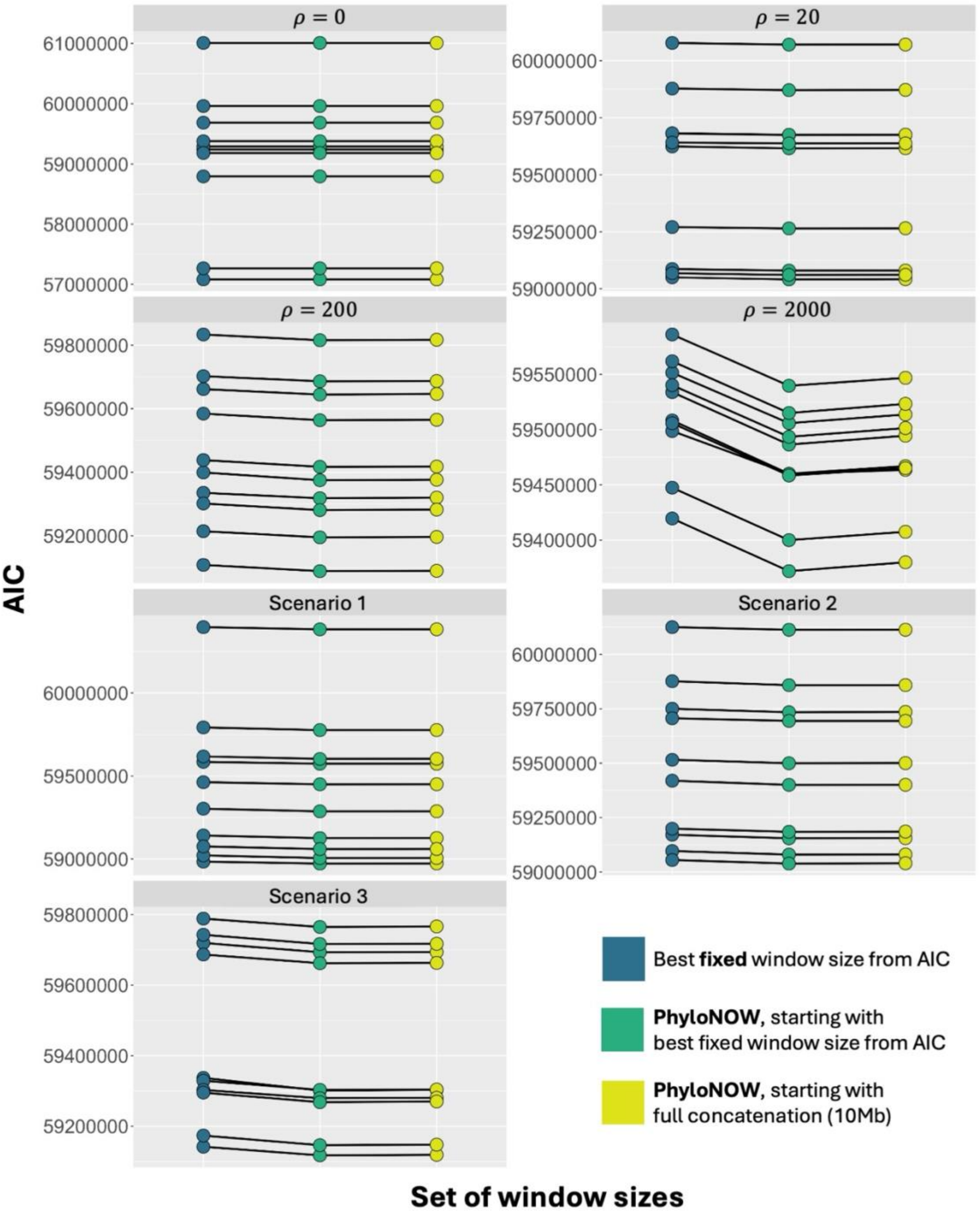


***Figure S1.*** *AIC from running the non-overlapping window analyses with fixed and variable window sizes across 7 simulation scenarios (Fig. 2). Colour denotes the set of window sizes used when running the non-overlapping window method, either using a fixed window size (Ivan et al. 2025) or PhyloNOW with different starting window sizes. Black lines connect the same replicates.*

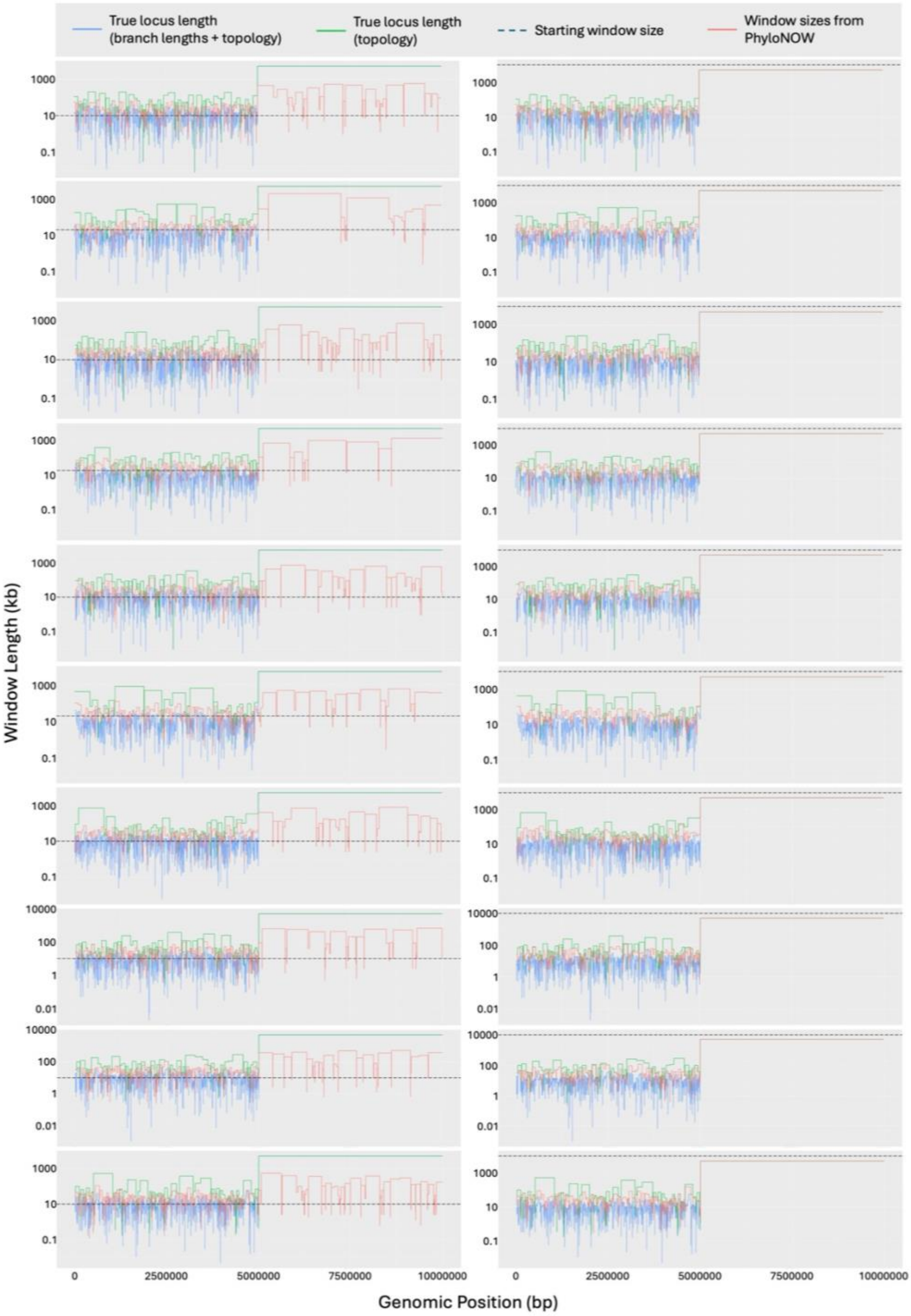


***Figure S2.*** *True simulated locus lengths and estimated window sizes from non-overlapping window analyses across simulated chromosomes from scenario 1 (Fig. 2). Blue lines: true locus lengths based on the branch lengths and tree topology; green lines: true locus lengths based on the tree topology only; black dashed lines: starting window size of the best fixed window size selected by the AIC (left) or 10Mb (right); red lines: window sizes from PhyloNOW with minimum 11 parsimony informative sites per window.*

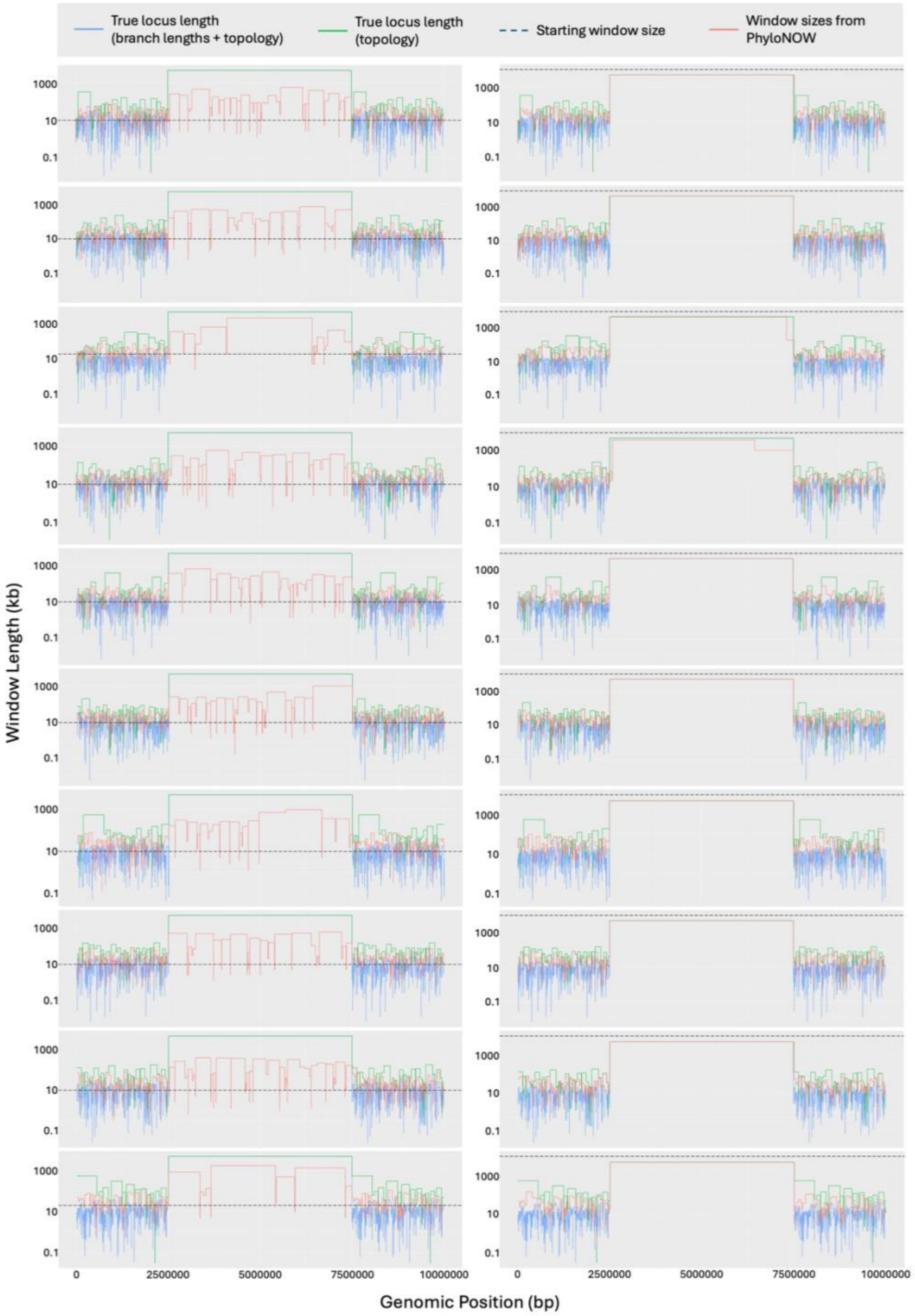


***Figure S3.*** *True simulated locus lengths and estimated window sizes from non-overlapping window analyses across simulated chromosomes from scenario 2 (Fig. 2). Blue lines: true locus lengths based on the branch lengths and tree topology; green lines: true locus lengths based on the tree topology only; black dashed lines: starting window size of the best fixed window size selected by the AIC (left) or 10Mb (right); red lines: window sizes from PhyloNOW with minimum 11 parsimony informative sites per window.*

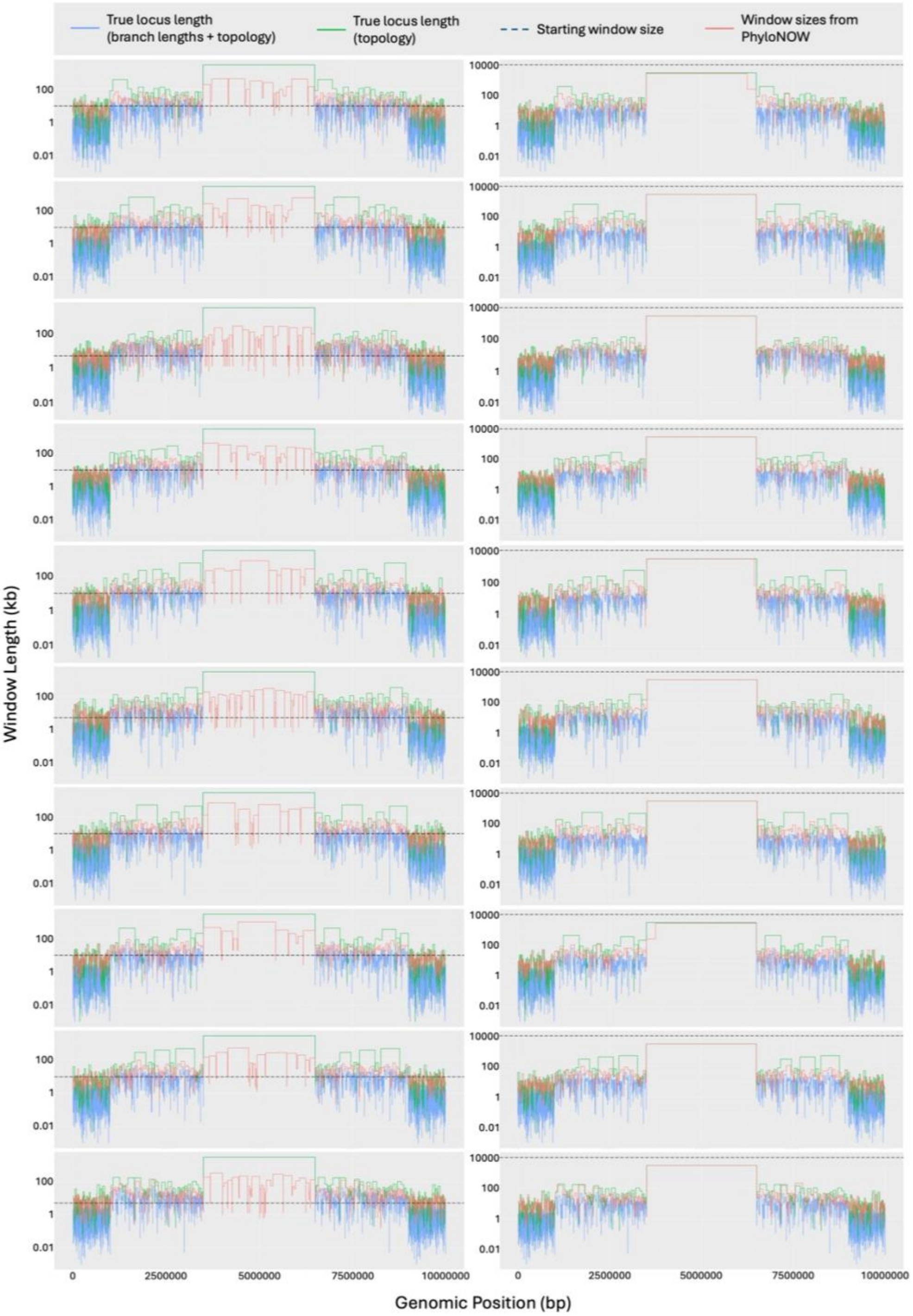


***Figure S4.*** *True simulated locus lengths and estimated window sizes from non-overlapping window analyses across simulated chromosomes from scenario 3 (Fig. 2). Blue lines: true locus lengths based on the branch lengths and tree topology; green lines: true locus lengths based on the tree topology only; black dashed lines: starting window size of the best fixed window size selected by the AIC (left) or 10Mb (right); red lines: window sizes from PhyloNOW with minimum 11 parsimony informative sites per window.*

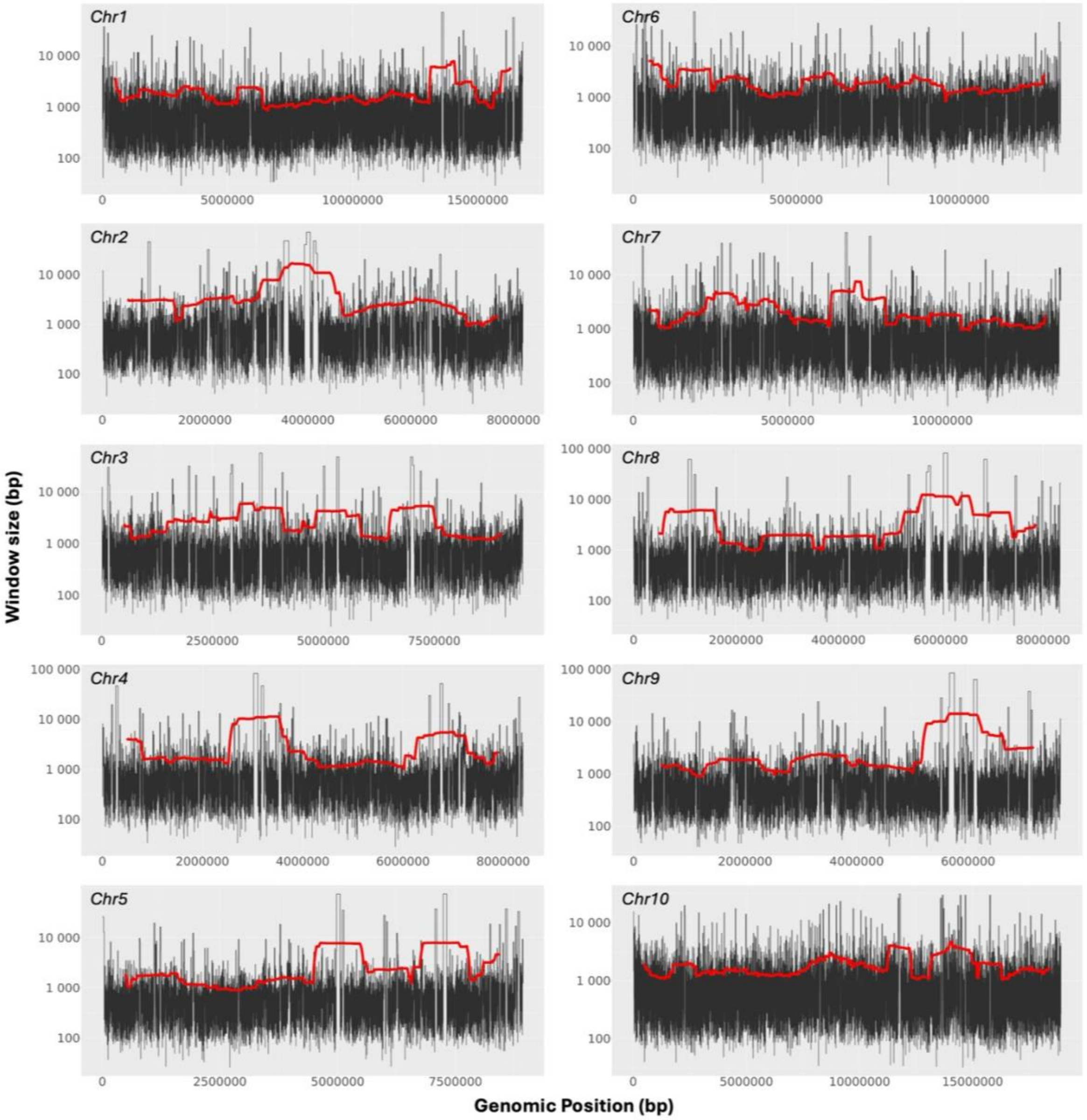


***Figure S5.*** *Variation of window sizes across the chromosomes of erato-sara clade of Heliconius butterflies. Black lines show the window size in base pairs (bp); red lines show the 1Mb moving average of the window sizes.*

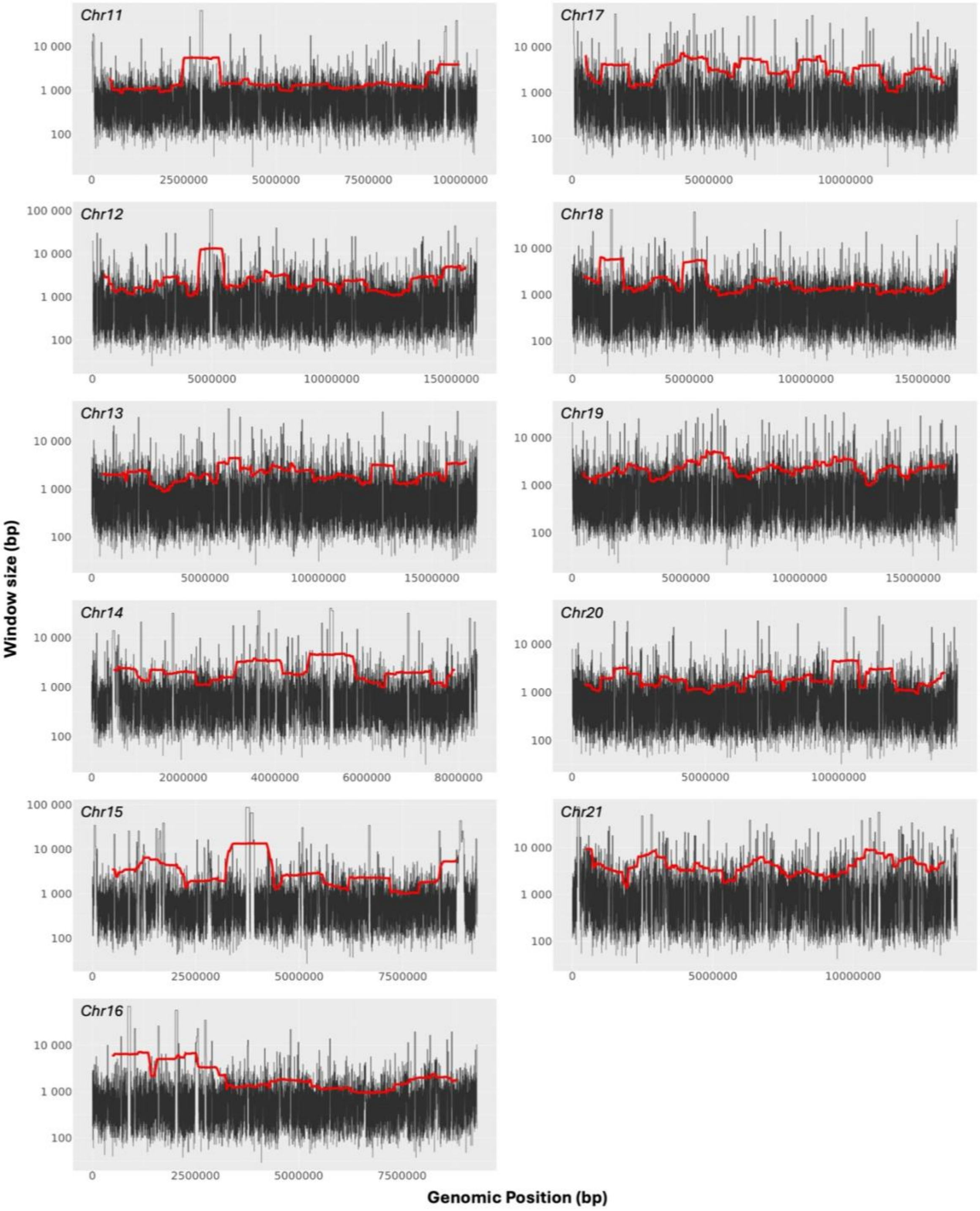


***Figure S5 (cont.).*** *Variation of window sizes across the chromosomes of erato-sara clade of Heliconius butterflies. Black lines show the window size in base pairs (bp); red lines show the 1Mb moving average of the window sizes.*

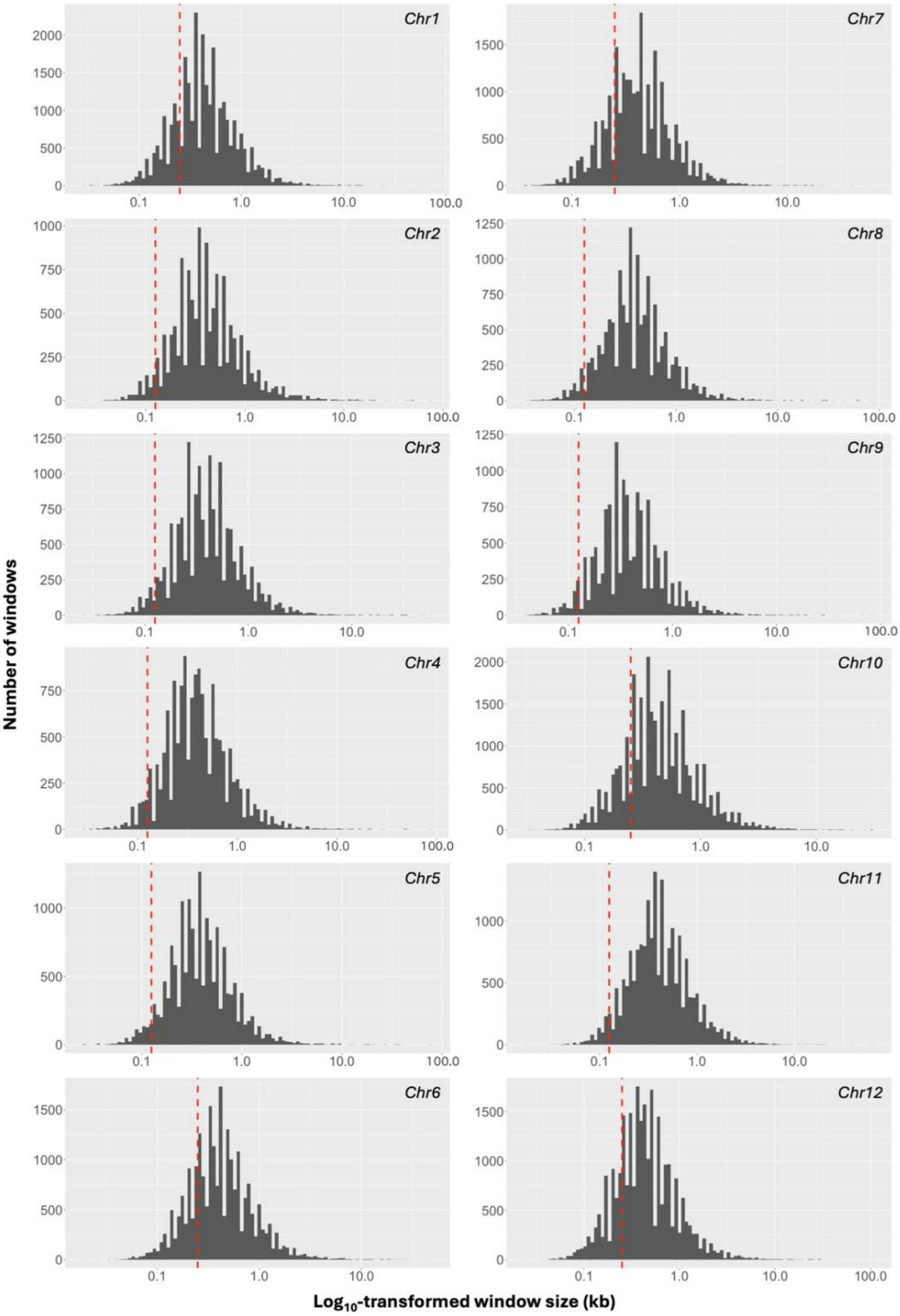


***Figure S6.*** *Distribution of window sizes across the chromosomes of erato-sara clade of Heliconius butterflies. Dashed red line denotes the best fixed window size based on the stepwise non-overlapping window analysis (Ivan et al. 2025).*

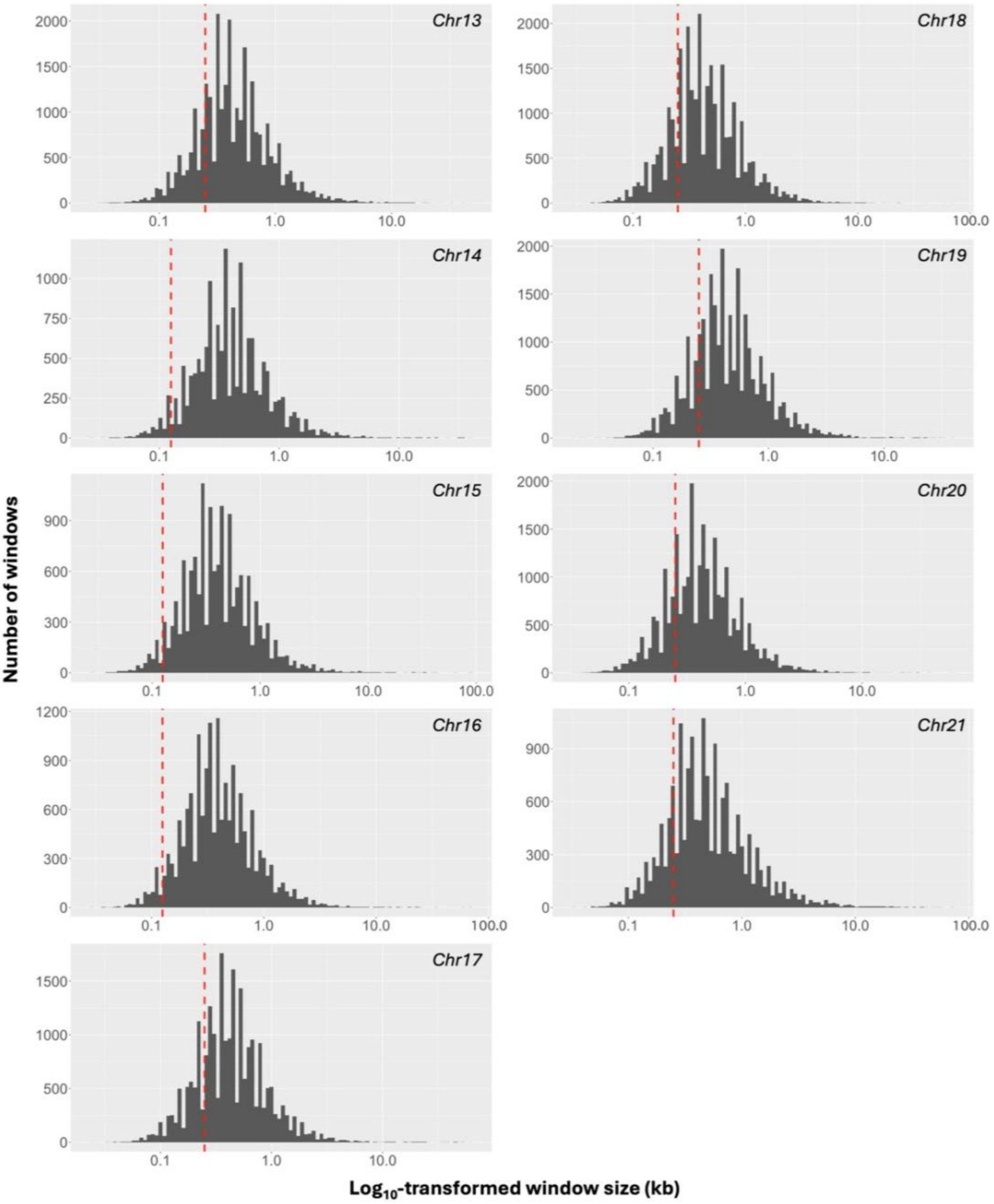


***Figure S6 (cont.).*** *Distribution of window sizes across the chromosomes of erato-sara clade of Heliconius butterflies. Dashed red line denotes the best fixed window size based on the stepwise non-overlapping window analysis (Ivan et al. 2025).*

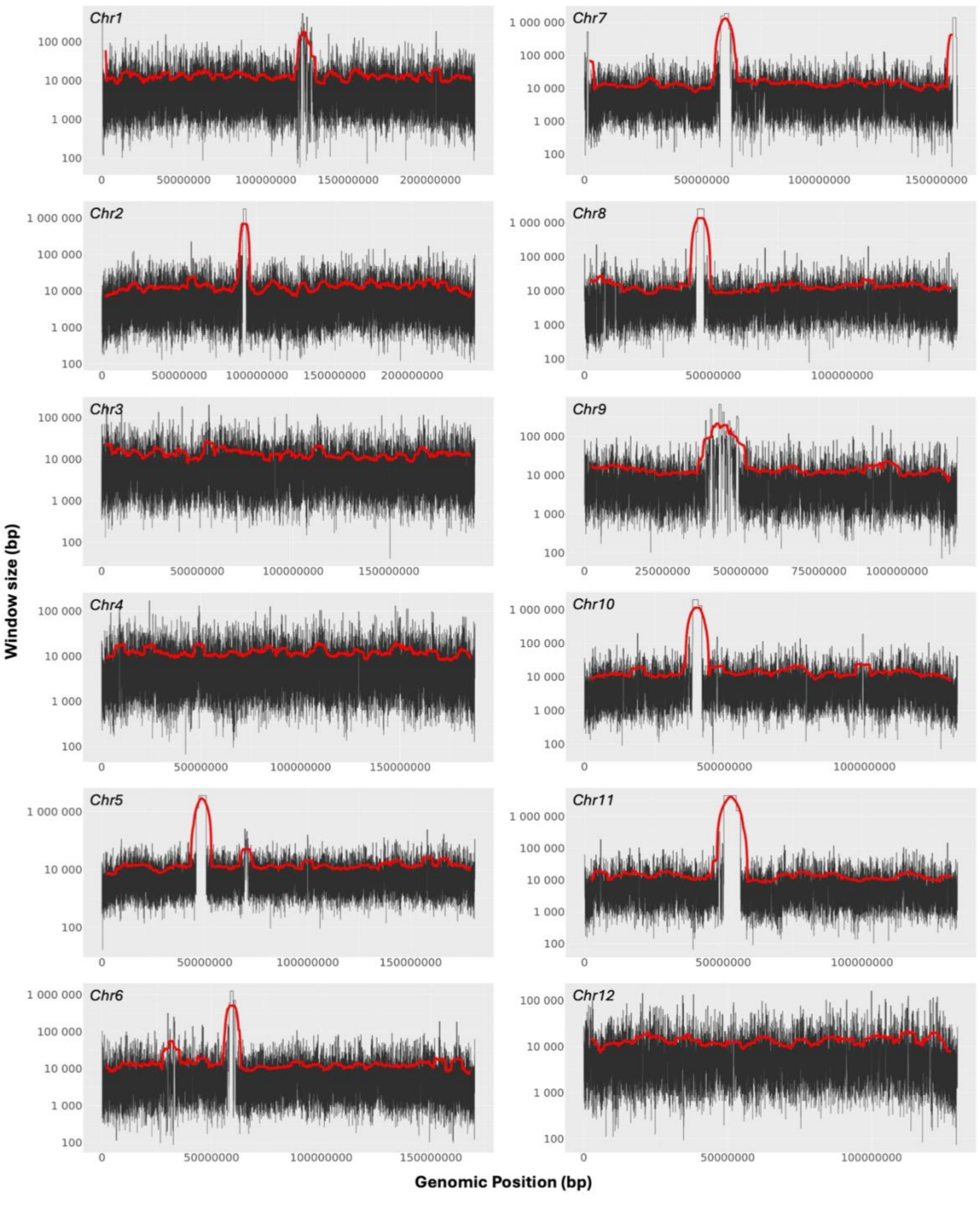


***Figure S7.*** *Distribution of window sizes across the chromosomes of great apes. Black lines show the window size in base pairs (bp); red lines show the 5Mb moving average of the window sizes.*

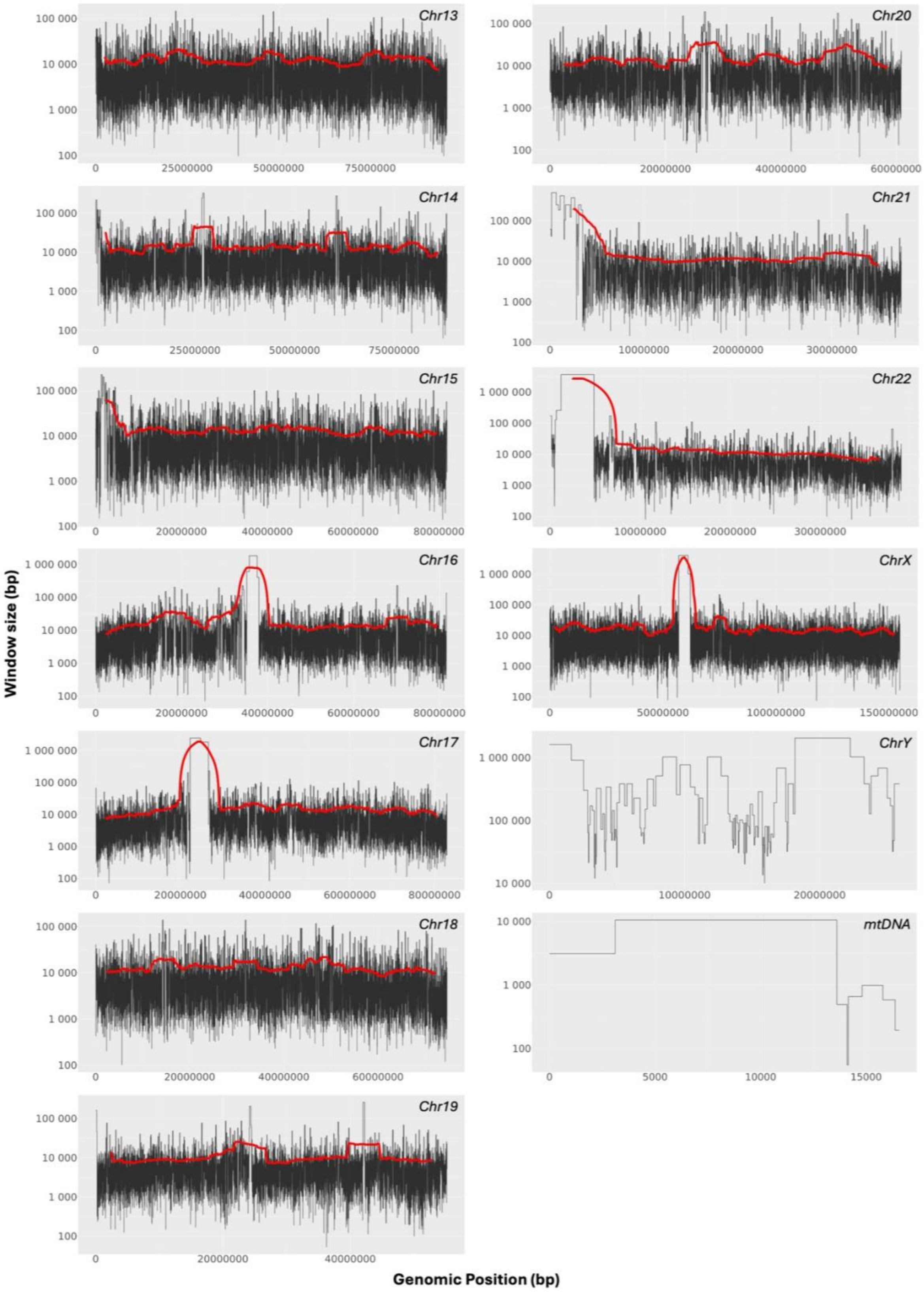


***Figure S7 (cont.).*** *Distribution of window sizes across the chromosomes of great apes. Black lines show the window size in base pairs (bp); red lines show the 5Mb moving average of the window sizes (except for chromosome Y and the mitochondrial genome).*

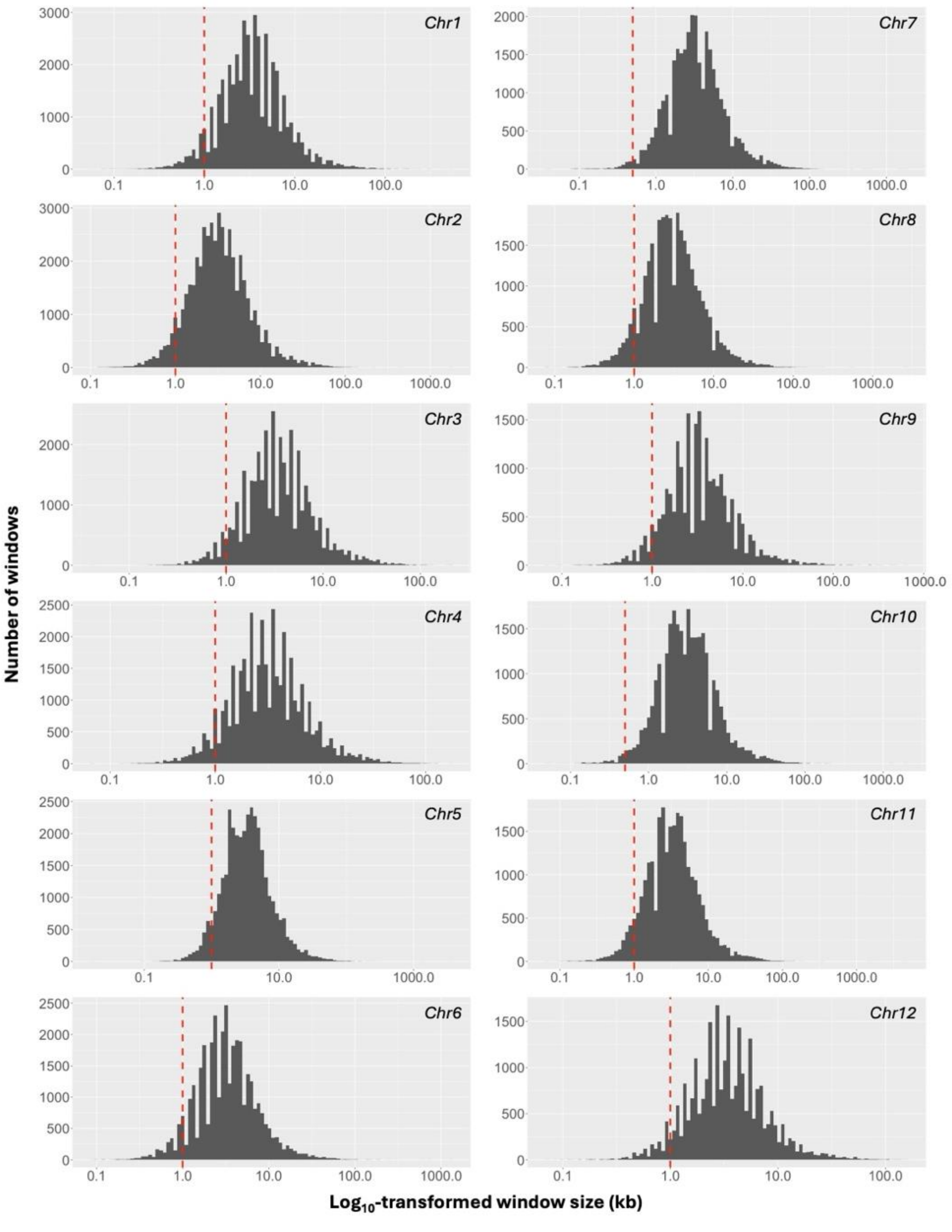


***Figure S8.*** *Distribution of window sizes across the chromosomes of great apes. Dashed red line denotes the best fixed window size based on the stepwise non-overlapping window analysis (Ivan et al. 2025).*

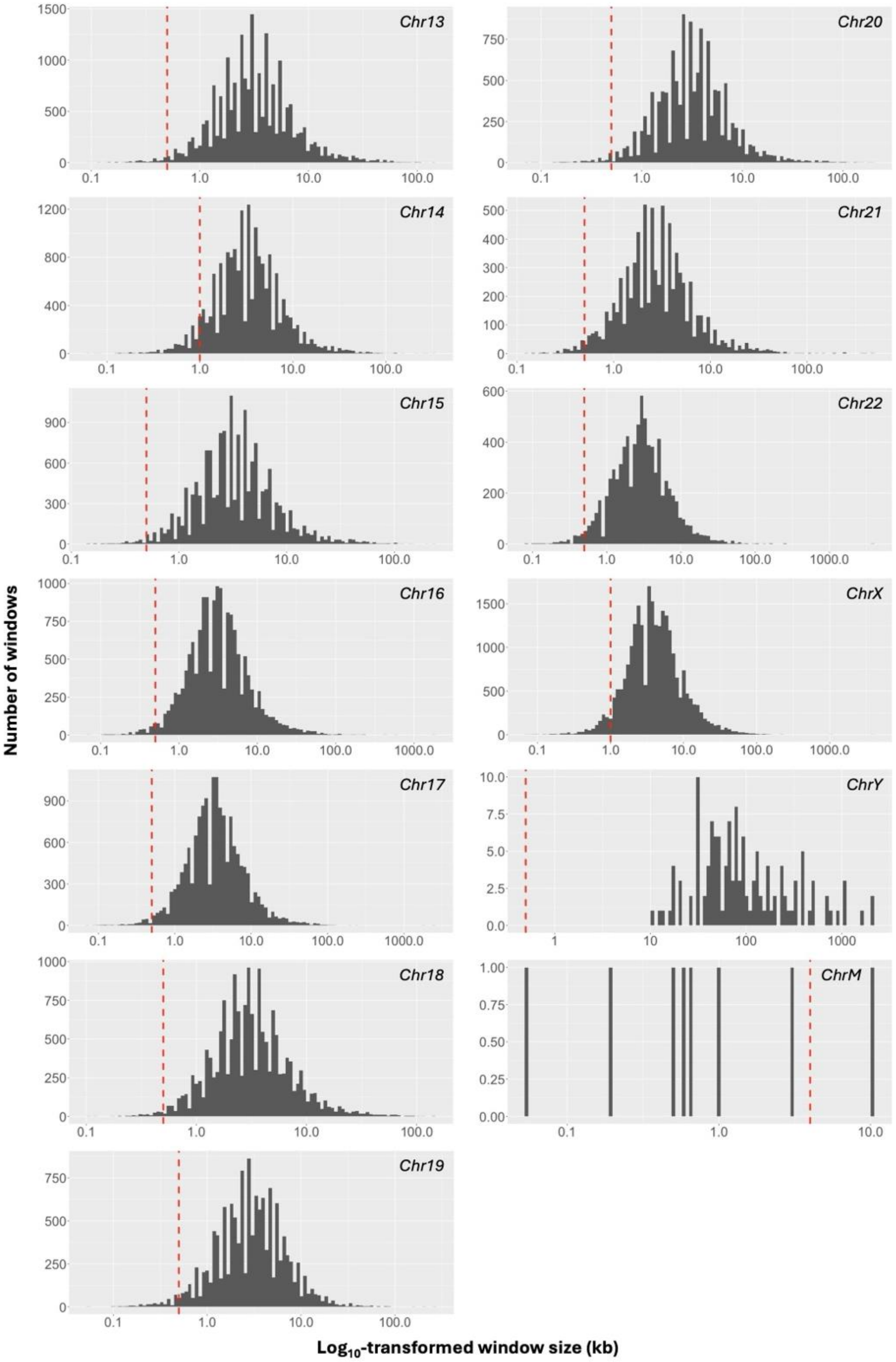


***Figure S8 (cont.).*** *Distribution of window sizes across the chromosomes of great apes. Dashed red line denotes the best fixed window size based on the stepwise non-overlapping window analysis (Ivan et al. 2025).*

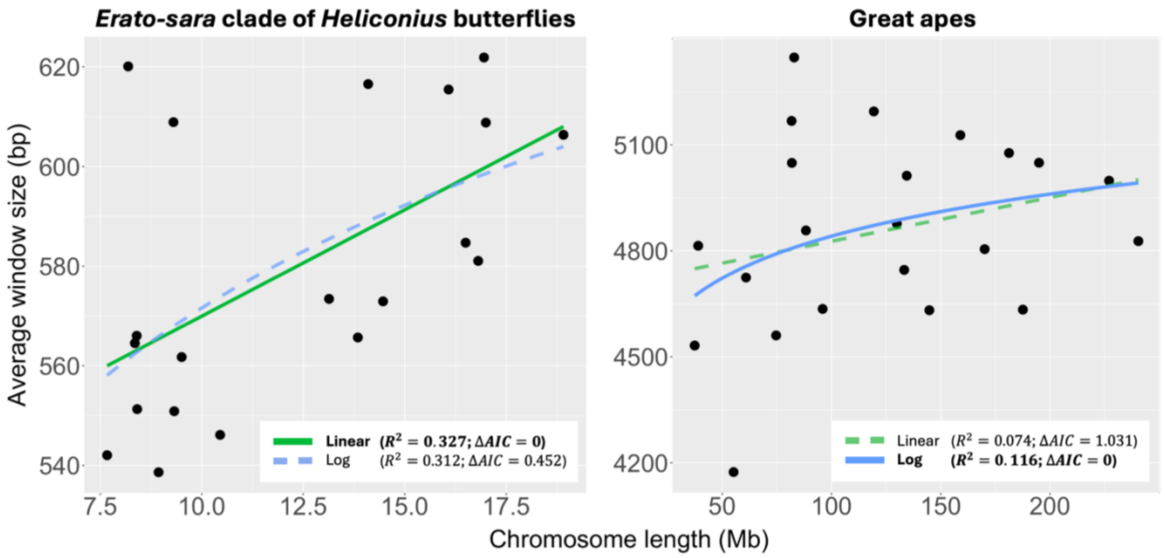


***Figure S9.*** *Correlation between chromosome lengths with average window sizes across autosomal chromosomes of erato-sara clade of Heliconius butterflies (left) and great apes (right). Colouring denotes different statistical models (i.e., linear, log), excluding asymptotic model as the model fails to converge. Model with the best AIC score is highlighted as solid line, while others are shown as dashed lines. ΔAIC reflects the AIC increase compared to the AIC of the best model.*

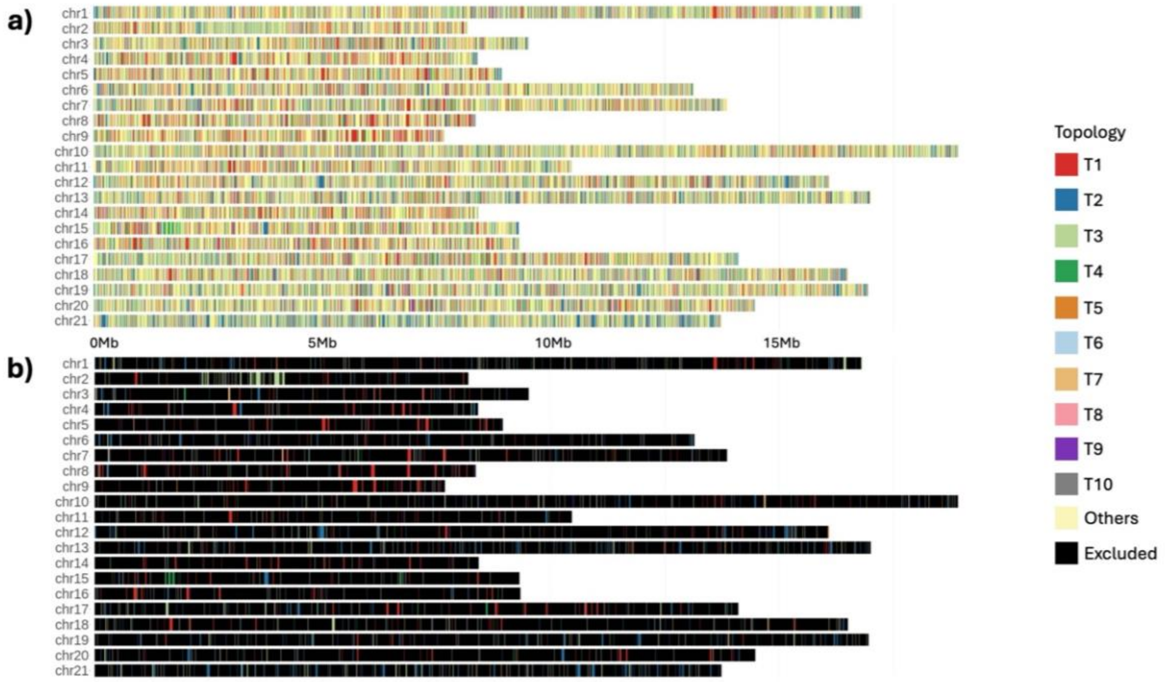


***Figure S10.*** *Topology distribution from the genomes of erato-sara clade of Heliconius butterflies based on (a) all gene trees and (b) gene trees with ≥95 average UFBoot support.*

*Colouring is based on Fig. 6, with Others for other topologies and Excluded for windows that are excluded from the analyses.*

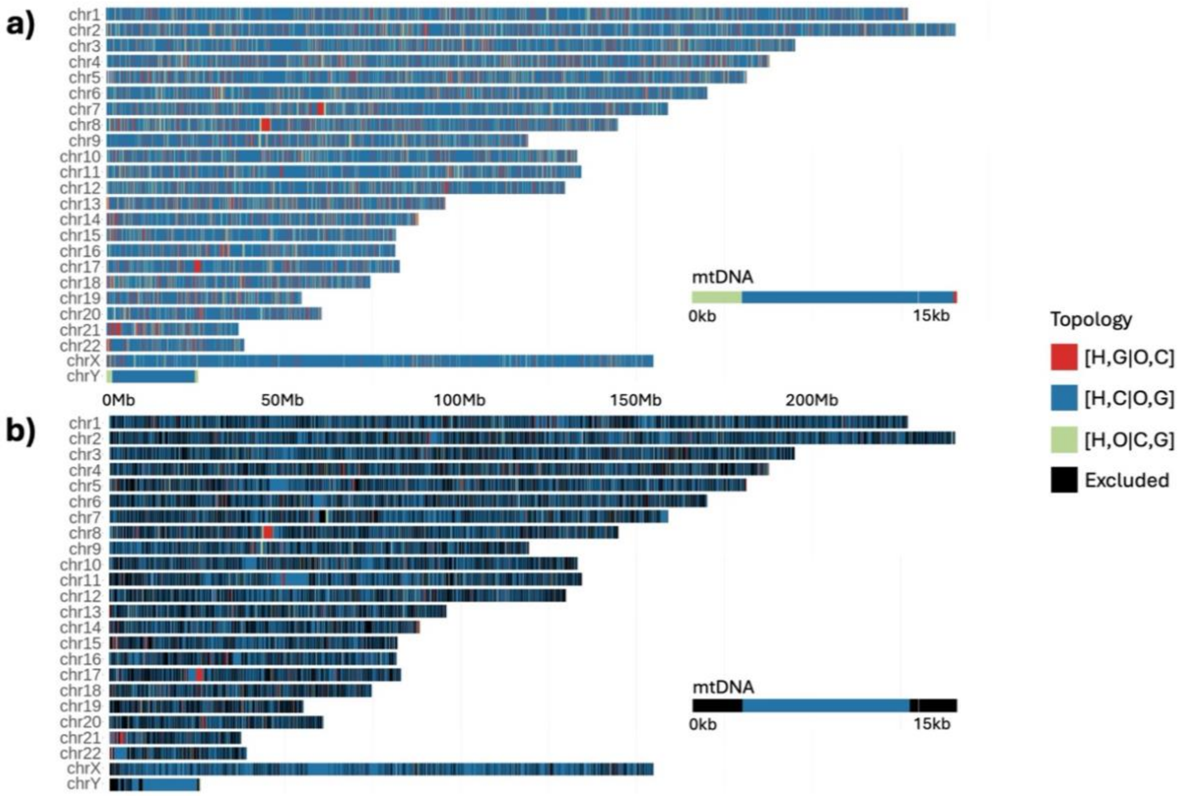


***Figure S11.*** *Topology distribution from the genomes of great apes based on (a) all gene trees and (b) gene trees with* $\geq$*95 average UFBoot support. Colouring is based on Fig. 7, with Others for other topologies and Excluded for windows that are excluded from the analyses. H: human; O: orangutan; G: gorilla; C: chimpanzee.*

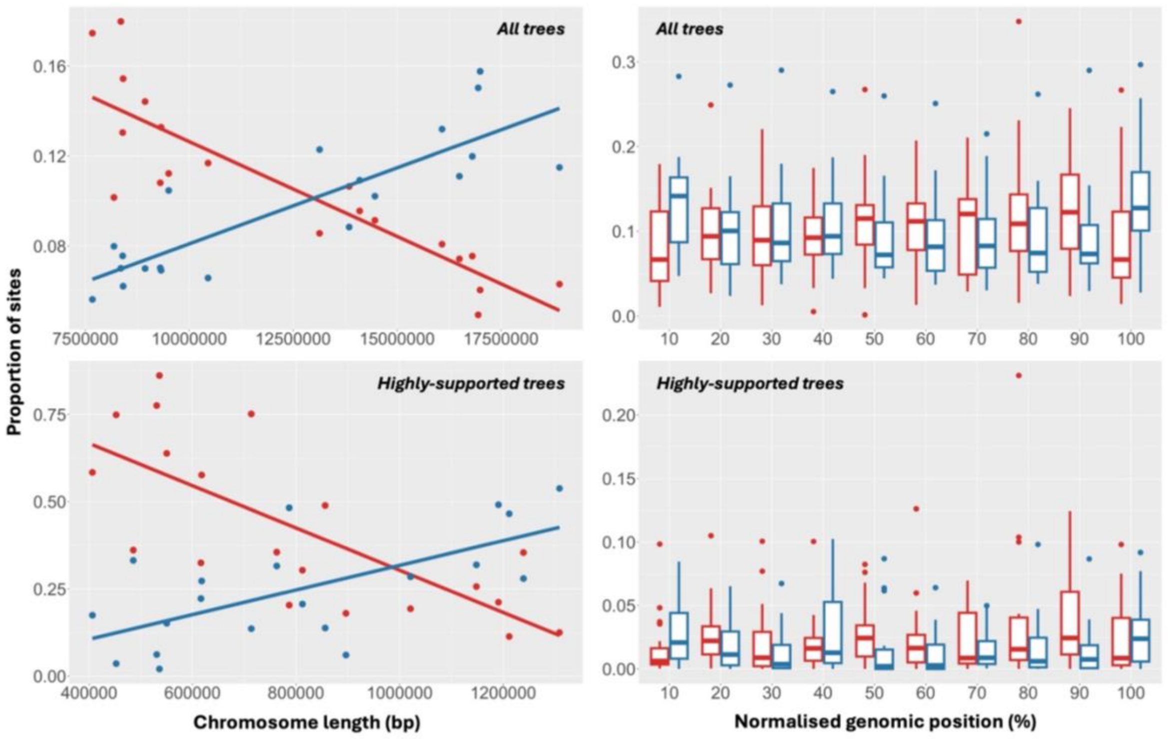


***Figure S12.*** *Correlation between the proportion of sites that recovered T1 (red) and T2 (blue) with the chromosomal architecture from erato-sara clade of Heliconius butterflies based on all gene trees (top) and gene trees with $\geq$95 average UFBoot support (bottom). Each dot denotes individual chromosome, except for chromosome 21.*
